# Combination of ALK2 and cholesterol targeting agents exploits linked genetic and metabolic dependencies in diffuse midline glioma

**DOI:** 10.64898/2026.09.23.753440

**Authors:** Rebecca Rogers, Alan Mackay, Yura Grabovska, Diana Carvalho, Claire Dobinson, Anna Burford, Valeria Molinari, Rita Pereira, Laura Bevington, Ruth Ruddle, Josephine Nkrumah, Haider Tari, Jiin Song, Drenusha Sejdiu, Nicole Chew, Vanessa Tsui, Claire Sun, Giulia Pericoli, Oren Becher, Maria Vinci, Jerome Fortin, Ron Firestein, Peter Sampson, Chris Jones

**Affiliations:** Centre for Children and Young People’s Cancer, Division of Cancer Biology, Institute of Cancer Research, London UK; Centre for Cancer Research, Hudson Institute of Medical Research, Monash University, Clayton, VIC 3168, Australia; Department of Molecular and Translational Science, Faculty of Medicine, Nursing and Health Sciences, Monash University, Clayton, VIC 3168, Australia; Research Area of Onco-haematology and Pharmaceutical GMP Facility, Bambino Gesù Children’s Hospital-IRCCS, Rome, Italy; Department of Pediatrics, Division of Pediatric Hematology-Oncology, the Mount Sinai Hospital Kravis Children’s hospital, New York, NY, USA; Department of Neurology and Neurosurgery, Montreal Neurological Institute-Hospital, McGill University, Montreal, Canada; Agora Open Science Trust, Toronto, Canada. M4K Pharma, Toronto, Canada

## Abstract

Diffuse midline glioma is an epigenetically driven disease defined by alterations targeting the histone post-translational modification H3K27me3 resulting in epigenetic rewiring and imposing unique metabolic dependencies. *ACVR1*-mutations arise in ∼25% of DMG H3K27-altered patients and impart a selective dependency on the kinase it encodes (ALK2), however ALK2 inhibitors show modest single-agent efficacy *in vivo.* In an attempt to better understand the cellular consequences of ALK2 inhibition to identify mechanistically-driven drug combinations, integrated multi-omics analysis was performed and revealed a novel role for ALK2 in cholesterol homeostasis while CRISPR and high-throughput drug screens identified hits targeting cholesterol metabolism as sensitisers to ALK2i. *In vivo* assessment of ALK2i plus clinically well-tolerated statins revealed a significant increase in the median survival compared to vehicle. Forced differentiation of DMG cells from an oligodendrocyte precursor-like to an astrocyte-like cell-state and led to a significant decrease in ALK2i sensitivity and synergy with statins, which was phenocopied when DMG cells were co-cultured with normal astrocytes. We identify a previously unappreciated role for ALK2 signalling in cholesterol homeostasis, showing cell-state dependency, and identify a rational combinatorial strategy for clinical translation.

**Significance:** Multi-omic characterisation of ALK2i response in *ACVR1*-mutant patient derived models of diffuse midline glioma reveals a linked metabolic and genetic dependency which can be exploited therapeutically by combining ALK2 inhibitors with clinically well-tolerated statins.

**Graphical abstract:** 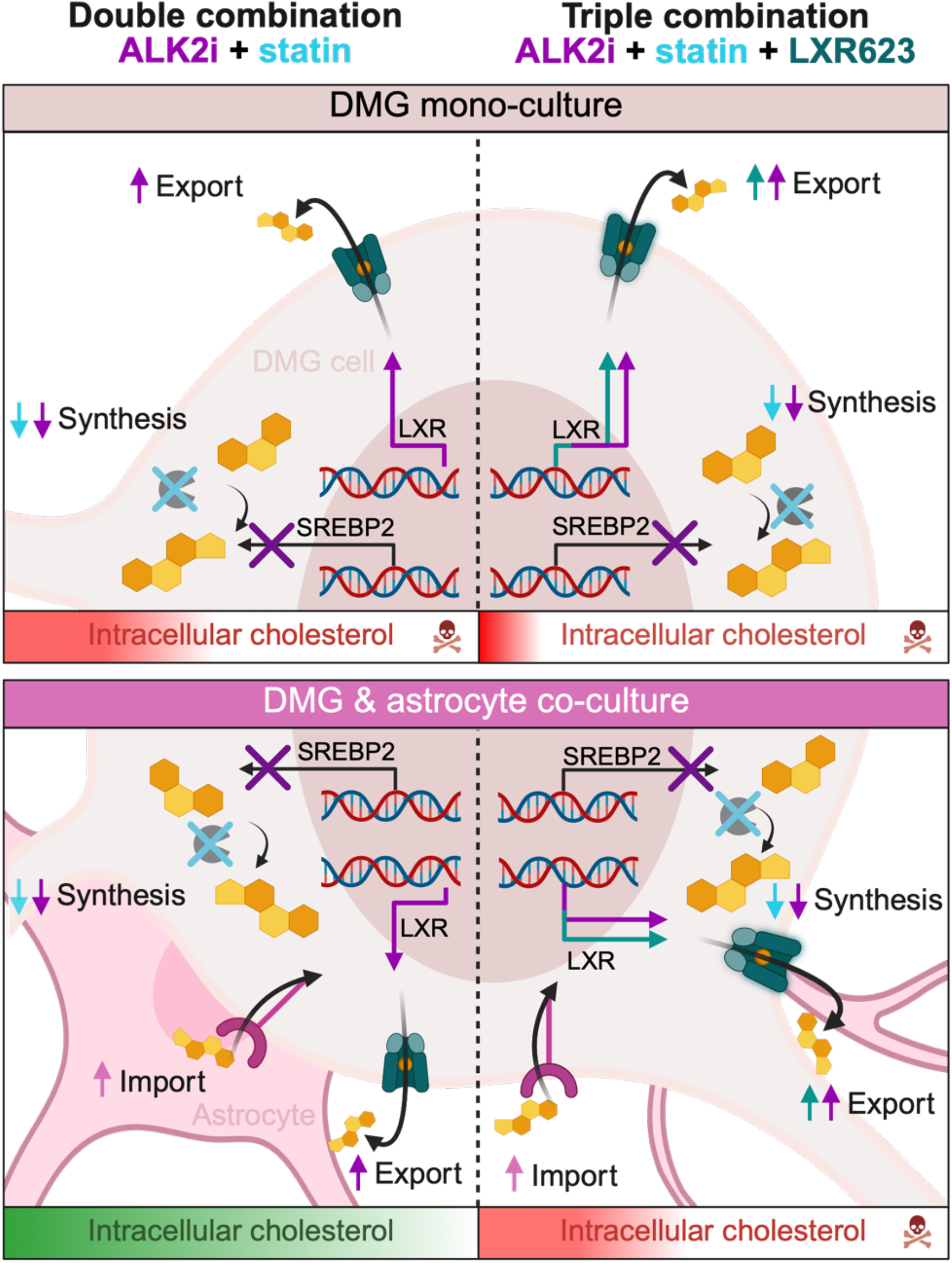

## Introduction

Diffuse midline glioma, H3K27-altered (DMG-H3K27), including diffuse intrinsic pontine glioma (DIPG), is a universally fatal diagnosis with a median survival of 9-15 months, an unmet need that has remained unchanged for decades [1–3]. DMG is an epigenetically driven disease defined by alterations targeting the histone post-translational modification H3K27me3, either by lysine-to-methionine substitutions at the lysine-27 residue itself, or via overexpression of the polycomb repressive complex 2 (PRC2) interactor EZHIP. The H3K27M mutation prevents PRC2 mediated deposition of H3K27me3 which leads to dysregulation and global changes of this largely repressive epigenetic mark [4–8]. This dysregulation alters gene expression and drives a stalled developmental cancer stem cell-like phenotype which closely resembles the oligodendrocyte precursor cell (OPC)-like state [9–12]. This epigenetic rewiring has also imposed unique metabolic dependencies which play a crucial role in DMG proliferation, including glycolysis, cholesterol metabolism and oxidative phosphorylation [13–17].

The somatic *ACVR1* mutations that arise in 20-25% DMG-H3K27-altered tumours, predominantly those with mutations in H3.1, represent an as yet unexploited target in DMG [18–21]. The identification of the gain of function mutations in *ACVR1,* which encodes the serine/threonine kinase ALK2, has driven the development of ALK2 inhibitors for targeted treatment. These include the potent tool compound LDN193189, the pre-clinical development compounds M4K2009 and M4K2163 and the clinical candidate Itacnosertib (TP-0184), which modulate the canonical ALK2 mediated BMP signalling pathway with varying potency and selectivity [22–27]. Despite substantial efforts to develop selective and potent ALK2 inhibitors [28] they have yet to be translated into the clinic, which is in part due to their modest single-agent efficacy *in vivo* with ∼10-15% increase in the median survival and all mice succumbing to disease [29–32]. The drug combination of vandetanib, (VEGFR/RET/EGFR inhibitor with off-target effects on ALK2) and everolimus (mTOR inhibitor) was identified using an AI-approach with a focus of drug repurposing and evaluated *in vivo*, which resulted in a similar extension of survival in pre-clinical models [31]. A better characterisation and understanding of ALK2 inhibitor treatment response and new combinatorial approaches are desperately needed in order to advance these inhibitors clinically.

In this study, using both multi-omic and high-throughput screening approaches, we evaluate the consequences of ALK2 inhibition in multiple *ACVR1*-mutant patient-derived models of DMG-H3K27. This reveals a novel role for ALK2 in cholesterol homeostasis and suggests a readily translatable combination approach with selective ALK2 inhibitors, taking advantage of tumour cell intrinsic and extrinsic mechanisms in the context of *ACVR1*-mutant DMG-H3K27.

## Results

### Drug-on CRISPR screens reveal loss of cholesterol homeostasis genes as sensitisers to ALK2 inhibitors

To identify novel combination partners which may potentiate ALK2 inhibitor potency in DMG, multiple drug-on combination CRISPR screens were carried out using sub-lethal concentrations of four compounds of different chemotypes across four patient-derived *ACVR1*-mutant DMG models, each harbouring different *ACVR1* mutations (Fig. 1a). The ALK2 inhibitors used included the clinical candidate TP-0184 [25], the drug development candidates M4K2009 and M4K2163 [24], as well as the tool compound LDN193189 [22, 23], each exhibiting distinct selectivity and potency for ALK2 (Extended data Fig.1a and Supplementary Table 1) [33]. We calculated the Δ Z-score (Z-score control – Z-score ALK2 inhibitor treatment) and subsequently ranked the differential hits for each screen (Fig. 1a and Supplementary Table 2). Within each institution the screens shared ∼ 6-10% of the top 500 differential hits (Fig. 1b-d and Supplementary Table 3), the overlap was lower when comparing between platforms and ALK2 inhibitors (Extended data Fig.1b), as evidenced by Jaccard similarity scores (Extended data Fig.1c). However, following Gene Set Enrichment Analysis (GSEA) analysis of the top-ranking ALK2 inhibitor sensitising hits, the degree pathway level agreement was highest between the more selective M4K compounds, irrespective of institute (Fig. 1e), revealing a convergence on cholesterol biosynthesis pathways highlighted by red text (Extended data Fig.1d-j). The top hits included *DHCR24* (24-dehydrocholesterol reductase), *EBP* (emopamil-binding protein) and *LSS* (lanosterol synthase) which all encode cholesterol synthesis enzymes functioning in the mevalonate pathway [34]. *SREBF2* (sterol regulatory element binding transcription factor 2) was also identified as a hit, encoding for the transcription factor SREBP2 which regulates a positive-feedback loop to maintain optimal intracellular cholesterol levels (Fig. 1f-g) [35]. Overall, these hits ranked more highly in the CRISPR screens run in combination with the more selective ALK2 inhibitors M4K2009/M4K2163 compared to the less selective TP-0184 (Fig. 1f).

**Figure 1:**
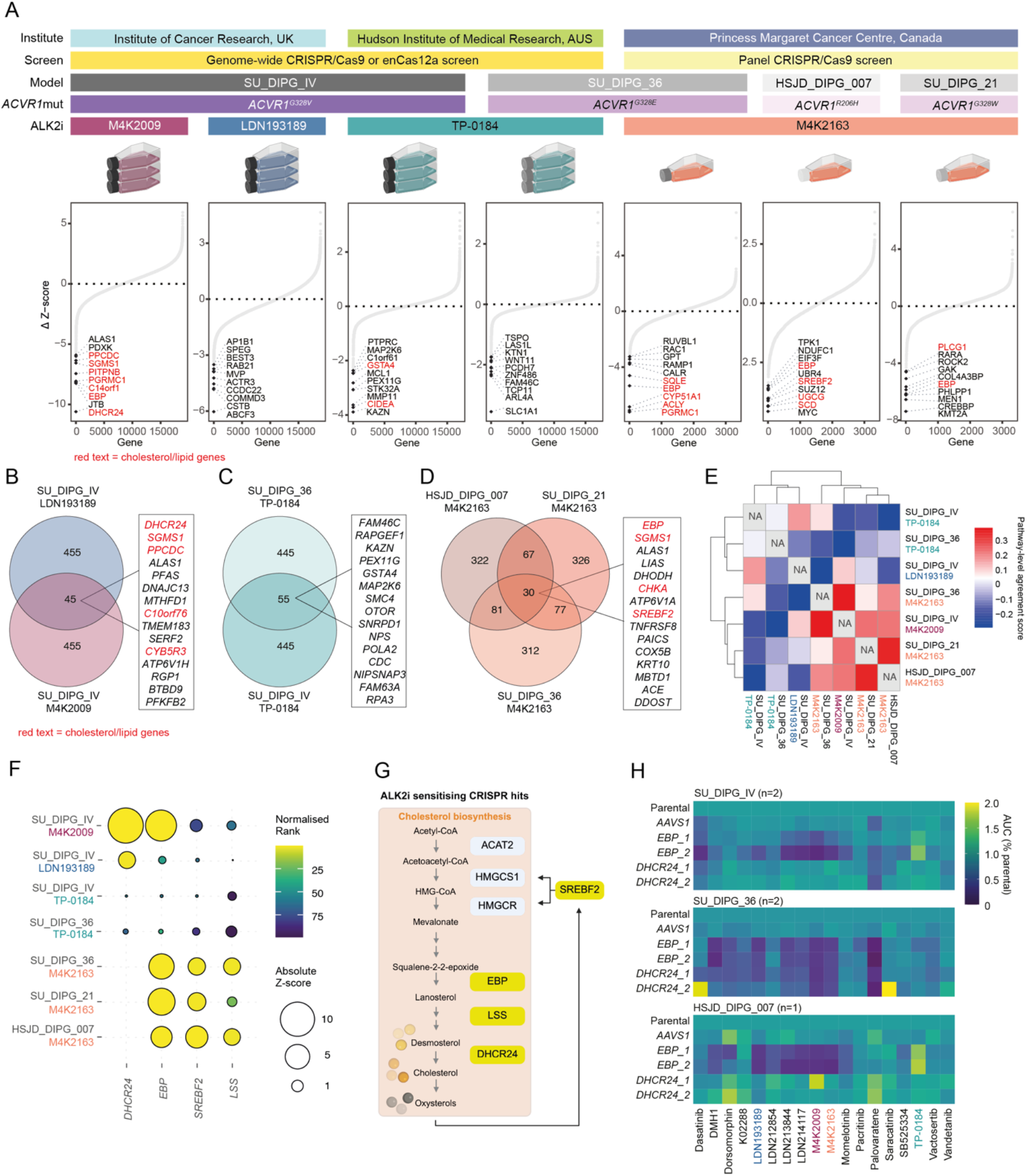
ALK2 inhibitor combination CRISPR screens reveal dependencies in cholesterol homeostasis**. A,** Waterfall plots of Δ Z-score (control Z-score – ALK2i Z-score, y-axis) for each gene (x-axis) for each of the CRISPR screens. The top 10 sensitising hits for each CRISPR screen have been labelled, red text indicates genes involved in cholesterol/lipid homeostasis. **B,** Venn diagram showing the overlapping ALK2 inhibitor sensitising hits (vehicle vs ALK2i) for CRISPR screens run at the Institute of Cancer Research. The top 15 overlapping genes are listed in the adjacent box, red text indicates genes involved in cholesterol/lipid homeostasis. **C,** Venn diagram showing the overlapping ALK2 inhibitor sensitising hits (vehicle vs ALK2i) for CRISPR screens run at the Hudson Institute for Medical Research. The top 15 overlapping genes are listed in the adjacent box, red text indicates genes involved in cholesterol/lipid homeostasis. **D,** Venn diagram showing the overlapping ALK2 inhibitor sensitising hits (vehicle vs ALK2i) for CRISPR screens run at the Princess Margaret Cancer Centre. The top 15 overlapping genes are listed in the adjacent box, red text indicates genes involved in cholesterol/lipid homeostasis. **E,** Heatmap showing the gene-set pathway-level agreement between CRISPR screens. Normalized enrichment scores (NES) were calculated for each pathway and experiment, and Pearson correlation of pathway NES profiles was used to quantify similarity between screens. The pathway-level agreement score is plotted by colour according to the key provided. **F,** Bubble plot showing the absolute Z-score (circle size) and normalised rank (according to colour key provided) of the prioritised hits (x-axis) across the six CRISPR screens (y-axis). **G,** Simplified cholesterol synthesis pathway schematic highlighting (yellow) the cholesterol synthesis enzymes encoded by the genes determined to be ‘hits’ in the CRISPR screens, above. **H,** Heatmap showing the AUC of a panel of ALK2i (fold-change to parental) in SU_DIPG_IV (top, n=2), SU_DIPG_36 (middle, n=2) and HSJD_DIPG_007 (bottom, n=1) following CRISPR-Cas9 directed gene knockout of prioritised hit genes, *AAVS1* negative gRNA control was used. The AUC (% parental condition) is plotted by colour according to the key provided.

Using an aggregated bulk RNAseq dataset of 320 paediatric-type diffuse high grade glioma (PDHGG) tumours, we observed significantly higher expression of the cholesterol homeostasis signature in the DMG-H3K27 subgroup compared to all other subgroups as defined by MNP12.8 (Extended data Fig.1k)[36]. Next, we sub-set the DMG-H3K27 tumours based on *ACVR1*-mutation status which revealed significantly higher expression of the cholesterol homeostasis signature in *ACVR1*-mutant tumours compared to *ACVR1*-wt (Extended data Fig.1l). Furthermore, differential expression analysis between *ACVR1*-mutant and *ACVR1*-wt H3K27-altered DMG and subsequent GSEA revealed a significant enrichment of cholesterol homeostasis, identifying a potentially novel role for *ACVR1*/ALK2 in cholesterol regulation (Extended data Fig.1m).

We prioritised *EBP* and *DHCR24* for validation and used both genetic and pharmacological tools to target these genes in three *ACVR1*-mutant patient-derived models. CRISPR/Cas9 knock-down of these genes alone minimally reduced the cell viability compared to the *AAVS1* control (Extended data Fig.1n), but increased sensitivity to multiple ALK2 inhibitors, most prominently with *EBP* knock-down, though this was not seen for TP-0184 (Fig. 1h).

### ALK2 inhibitors plus statins show strong synergy in PDHGG models

An orthogonal high-throughput drug screening approach was used to identify potential drug combination partners using ALK2 inhibitors with different selectivity profiles (M4K2009 or TP-0184) (Fig. 2a). Differential robust Z-scores for each compound (+/- ALK2i) were calculated (Supplementary Table 4) and pathway analysis of the top 30 hits revealed enrichments in compounds targeting neuronal signalling, cholesterol metabolism and hormone receptors (HR) for M4K2009 and PI3K/AKT/MTOR, tyrosine kinase signalling and metabolism for TP-0184 (Fig. 2b). Interestingly, the hormone receptor hits specifically associated with M4K2009 included multiple estrogen receptor inhibitors, including tamoxifen, which have reported off-target effects on EBP [37]. Common hits between the screens included inhibitors targeting cholesterol synthesis via HMG-CoA reductase (lovastatin and simvastatin [38]), the receptor tyrosine kinases VEGFR/FGFR/EGFR, and other cancer-related kinases such as ATR and PIM-kinase (Fig. 2c). Due to the convergence of both the CRISPR and drug screening hits on cholesterol homeostasis, we focused our validation on the clinically routinely-used statins. Screening a panel of patient-derived models, representing different PDHGG subgroups (Extended data Fig.2a and Supplementary Table 5), differential single-agent sensitivity to simvastatin and M4K2009 was observed compared to normal human astrocytes (HA-FL, HA-BS), neuronal stem cells (E3462) and oligodendrocytes (MO3.13) (Fig. 2d-e). Bliss combination assays highlighted robust synergistic interactions (>10) between statins and M4K2009 across a panel of patient-derived *ACVR1*-mutant models while only weak additivity/antagonism (<2) was observed in the normal lines tested (Fig. 2f and Extended data Fig.2b), suggesting a potential therapeutic window. We also observed synergy between statins and M4K2009 in a wider panel of histone-mutant PDHGG models (Extended data Fig.2c) suggesting that ALK2 inhibitor combinations might also be worth exploring in other subgroups of PDHGG.

**Figure 2:**
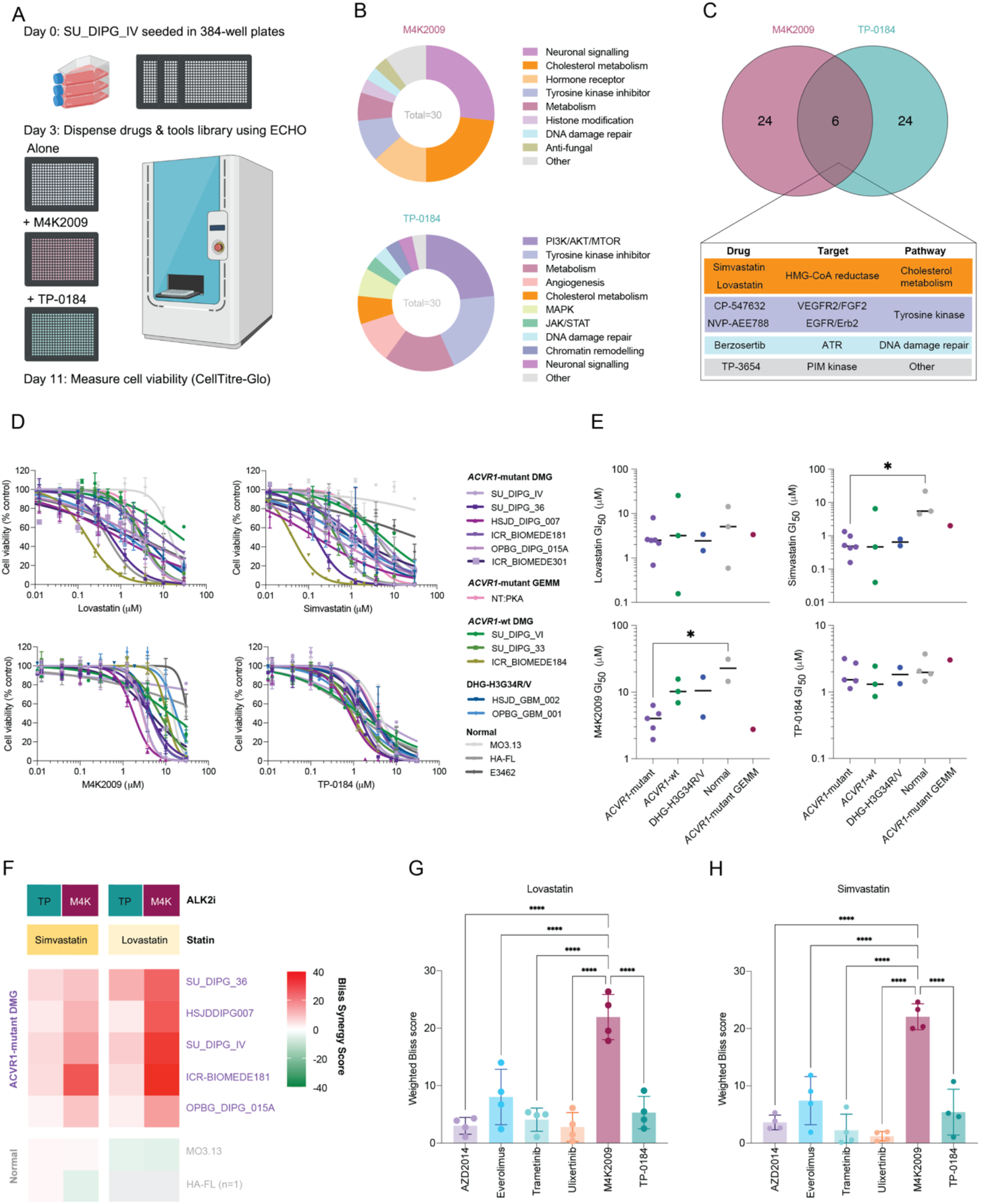
High-throughput ALK2 inhibitor drug combination screens identify cholesterol targeting inhibitors as top overlapping hits. **A,** Schematic of high-throughput ALK2 inhibitor (M4K2009 or TP-1084) combination screens performed in 384-well plates using SU_DIPG_IV cells. **B,** Donut plots showing the proportion of the different drug classes represented by the top 3% ALK2i sensitising hits identified from the M4K2009 (top) or TP-0184 (bottom) drug combination screens (n≥3). **C,** Venn of the overlapping ALK2i sensitising drug hits (top 3%) between M4K2009 and TP-0184. The drug targets and pathway details for the overlapping hits are detailed in the box below. **D,** Concentration-response curves for statins and ALK2 inhibitors across a panel of patient-derived PDHGG models and non-PDHGG models (n≥3). Data normalised to DMSO vehicle control, data shown as mean ± SD. Curves were fitted using [inhibitor] vs response, variable slope, four parameters in Prism. **E,** GI_50_ values determined for statins and ALK2 inhibitors across a panel of patient-derived PDHGG models and non-PDHGG models (MO3.13, HA-FL and E3462) split by subgroup / genotype (n≥3). p values calculated using one-way Anova, *= p<0.05. **F,** Heatmap of Bliss synergy scores for M4K2009 (M4K) or TP-0184 (TP) in combination with statins in a panel of patient-derived *ACVR1*-mutant and non-PDHGG models (MO3.13 and HA-FL) (n=3). Bliss scores > 10 indicate synergy and <10 indicate antagonism and are plotted by colour according to the key provided. **G,** Barplot of weighted Bliss synergy scores for a panel of clinically relevant mTOR/ERK/MEK inhibitors or ALK2 inhibitors in combination with lovastatin in three patient-derived PDHGG models and a mouse *ex vivo* syngeneic model (RCAS model n=1, patient-derived modes n≥2). p values calculated by one-way ANOVA (****p<0.0001). **H,** Barplot of weighted Bliss synergy scores for a panel of clinically relevant mTOR/ERK/MEK inhibitors or ALK2 inhibitors in combination with simvastatin in three patient-derived PDHGG models and a mouse *ex vivo* syngeneic model (RCAS model n=1, patient-derived modes n≥2). p values calculated by one-way ANOVA (****p<0.0001).

The observed statin combination synergies with ALK2 inhibitors were benchmarked against other clinically relevant targeted agents, including inhibitors of the PI3K/mTOR and MAPK pathways [31, 39–41]. Although some synergy was observed, these combinations were mostly additive (mean weighted Bliss scores ranging from 1-8) and significantly lower than observed for M4K2009 plus statins (mean weighted Bliss score = 22) (Fig. 2g-h). This combination also significantly out-performed statin plus TP-0184 (mean weighted Bliss score = 5), the latter of which has a substantially lower selectivity for ALK2 over ALK5 (TGFBR1) than M4K2009 (7- vs 187-fold selectivity ALK2/ALK5, respectively)[42].

To assess the contribution of ALK2 and ALK5 activity to statin sensitivity we performed CRISPR/Cas9 knock-down experiments targeting either *ACVR1* (encoding ALK2) or *TGFBR1* (encoding ALK5) in SU_DIPG_IV cells and assessed knock-down efficiency using TIDE analysis (Extended data Fig.2d). *ACVR1* and *TGFBR1* knock-down alone led to a partial reduction in cell viability (Extended data Fig.2e), however only *ACVR1* knock-down resulted in sensitisation to statin treatment (Extended data Fig.2f-g). Together, these observations suggest the effects of ALK2 inhibitors on cholesterol homeostasis in the context of DMG, and the resultant high degree of synergy with cholesterol biosynthesis inhibitors is mediated by ALK2 itself, not off-target effects on ALK5.

### Multi-omic profiling reveals novel role for ALK2 in cholesterol homeostasis

To investigate a possible mechanistic link between ALK2 signalling and cholesterol metabolism, the global downstream consequences of ALK2 inhibition were evaluated at the transcriptomic, proteomic and metabolomic levels (Fig. 3a). Integrated analysis of the transcriptomics and proteomics showed that M4K2009 or TP-0184 treatment results in a down-regulation of the BMP signalling marker ID1, confirming an on-target effect of both inhibitors (Fig. 3b). The most differentially upregulated gene/protein for both compounds was the cholesterol exporter ABCA1, usually upregulated under high cholesterol conditions via a feedback loop triggered by oxysterol binding and activation of the transcription factor liver X receptor (LXR) (Fig. 3b)[43]. Immunofluorescent staining confirmed an increase in ABCA1 expression following 24h ALK2 inhibitor treatment (Extended data Fig.3a-b). The differentially expressed genes/proteins were subjected to GSEA and highlighted cholesterol synthesis and transport/export pathways as commonly depleted or enriched, respectively (Fig. 3c and Extended data Fig.3c).

**Figure 3:**
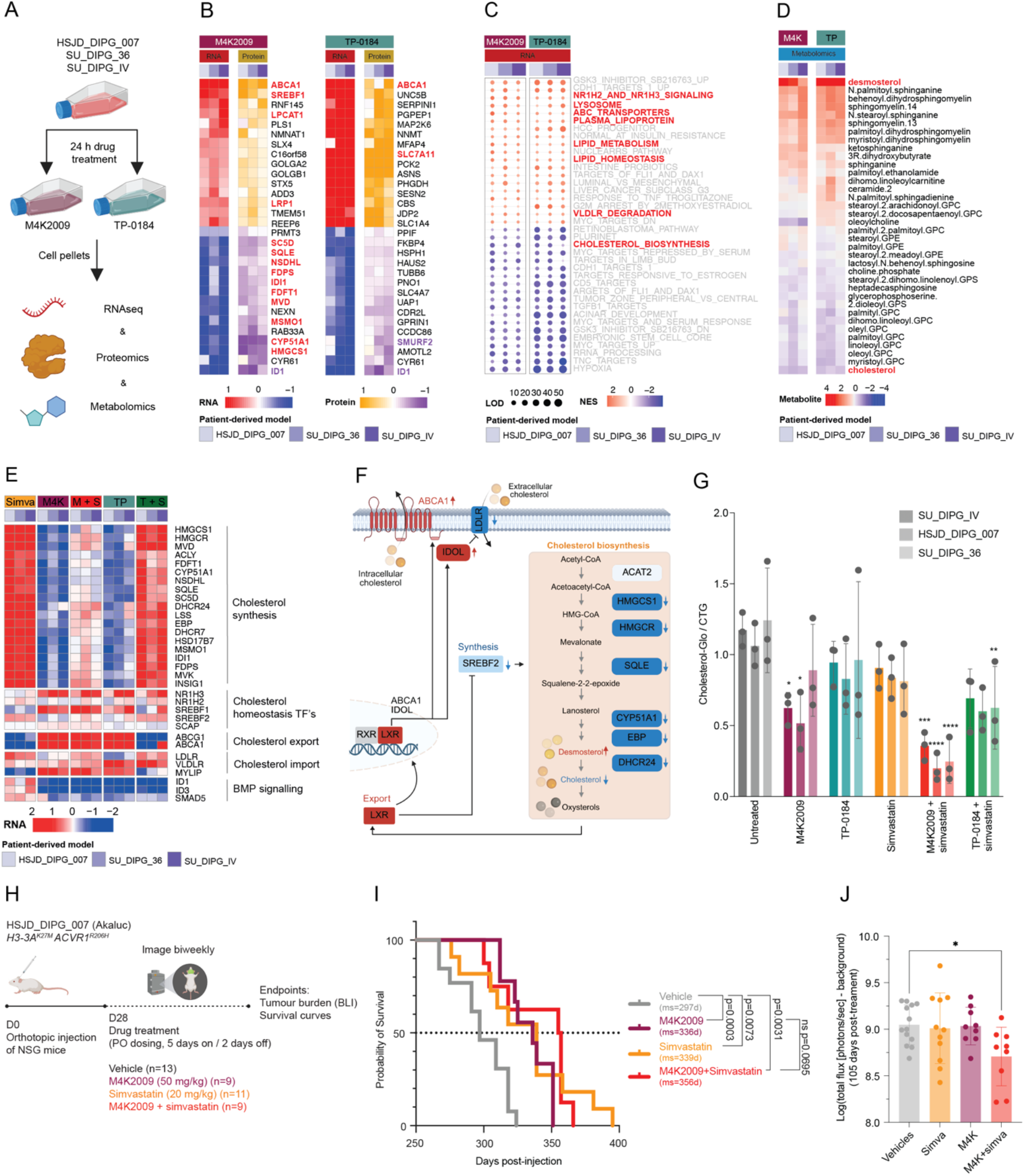
Multi-omic profiling of ALK2 inhibitor response in *ACVR1*-mutant patient-derived models reveals a novel role in cholesterol homeostasis. **A,** Schematic detailing the *ACVR1*-mutant patient-derived models, drug treatments and downstream multi-omic analysis performed. **B,** Heatmap of differentially expressed genes (red/blue) and proteins (yellow/purple) across three *ACVR1*-mutant patient-derived models following 24 h treatment with equipotent concentrations of M4K2009 (left) or TP-0184 (right). Data is mean of three biological replicates. Red text indicates genes involved in cholesterol/lipid homeostasis and purple text indicates those involved in BMP signalling. The expression is plotted by colour according to the key provided. **C,** Bubble plot of enriched or depleted pathways identified by GSEA of differentially expressed genes across three *ACVR1*-mutant patient-derived models. Red text indicates pathways associated with cholesterol/lipid homeostasis. The NES score is plotted by colour according to the key provided and the LOD is indicated by circle size. **D,** Heatmap showing the expression the lipid class metabolites in three *ACVR1*-mutant patient-derived models following 24 h treatment with equipotent concentrations of M4K2009 or TP-0184. Data normalised to control (DMSO) condition for each patient-derived model. Data is mean of five biological replicates. The expression is plotted by colour according to the key provided. **E,** Simplified cholesterol biosynthesis pathway schematic highlighting the cholesterol synthesis enzymes and metabolites that are up-(red) and down-regulated (blue) following 24 h ALK2i treatment. **F,** Heatmap showing gene expression associated with cholesterol homeostasis and BMP signalling across three *ACVR1*-mutant patient-derived models following 24 h of single-agent (M4K2009, TP-0184, simvastatin) or combinatorial treatment (ALK2 inhibitor + simvastatin). Data is mean of three biological replicates. The expression is plotted by colour according to the key provided. **G,** Cholesterol-Glo luminescence (y-axis) following 48 h of single-agent or combination treatment using equipotent concentrations in three *ACVR1*-mutant patient-derived models. Data normalised to CTG luminescence, shown as mean ± SD (n=3). p values calculated by two-way ANOVA (* p<0.05, ** p<0.01, *** p<0.001, **** p<0.0001) compared to untreated condition. **H,** Schematic of *in vivo* efficacy (survival) and tumour burden experiments using the orthotopic CDX model HSJD_DIPG_007-Akaluc in NSG mice. Oral dosing (PO) of vehicle, M4K2009 50 mg/kg, simvastatin 20 mg/kg or M4K2009 50 mg/kg and simvastatin 20 mg/kg was administered 5 days on, 2 days off. **I,** Survival analysis of single-agent (M4K2009 50 mg/kg, simvastatin 25 mg/kg) or combination treatment in tumour-bearing mice (HSJD_DIPG_007-Akaluc) using Log-rank (Mantel-Cox) test (vehicle vs simvastatin **p=0.0073, vehicle vs M4K2009 ***p=0.0003, vehicle vs M4K2009 plus simvastatin **p=0.0031, M4K2009 vs M4K2009 plus simvastatin ns p=0.0695). Mice were dosed via PO 5 days on/2 days off. Vehicle, n=13; simvastatin, n=11; M4K2009, n=9; M4K2009 plus simvastatin, n=9. Median survival (ms) in days is indicated in brackets. **J,** Bioluminescence intensity (BLI) was determined by IVIS imaging and used as a surrogate measure of tumour burden. Log total flux [photon/sec] minus background was calculated for each mouse and plotted per treatment arm; data shown is 105 days post-treatment. Data shown as mean ± SD. Dots represent the individual BLI signal for each mouse. p values calculated by one-way ANOVA (* p<0.05) compared to vehicle arm.

Global metabolomics revealed ALK2 inhibitor treatment leads to a raft of metabolomic changes, with the largest number of changes in the lipid class of metabolites (Fig. 3d, Extended data Fig.3d and Supplementary Table 6). Importantly, the expression changes we observed translated into a reduction in cellular cholesterol levels and increase in desmosterol (Fig. 3d), an immediate precursor to the synthesis of cholesterol (Bloch pathway) which is catalysed by DHCR24 (CRISPR screen hit) (Fig. 3f) [44]. Desmosterol has been shown to bind and upregulate LXR target genes including *ABCA1* [45, 46], here we show that *ACVR1*-mutant cells supplemented with exogenous desmosterol exhibit increased ABCA1 IF staining (Extended data Fig.3a-b). ALK2 inhibitor treatment also led to the reduction of Coenzyme A, a key metabolite and rate-limiting substrate in lipid/sterol biosynthesis (Extended data Fig.3e)[47]. The precursor metabolite of Coenzyme A (4’-phosphopantetheine) is synthesised by PPCDC; the *PPCDC* gene was identified as an ALK2 inhibitor sensitising CRISPR hit (Fig. 1a-b). Interestingly, the metabolite 3-hydroxy-3-methylglutarate was consistently increased following M4K2009 treatment and indicates a reduction in HMG-CoA metabolism [48], this was however more variable across the models following TP-0184 treatment (Extended data Fig.3f).

Despite M4K2009 and TP-0184 single-agent treatment similarly impacting cholesterol homeostasis, we consistently observed stronger synergy with statin plus M4K2009 treatment compared to statin plus TP-0184. To further investigate this, we assessed gene expression changes (by RNAseq) and cholesterol levels (using the CholesterolGlo assay) following combinatorial treatment of ALK2 inhibitor plus the HMG-CoA reductase inhibitor simvastatin. Strikingly, ALK2 inhibitor and simvastatin single-agent treatment resulted in opposing transcriptional responses with respect to cholesterol homeostasis (Fig. 3e), suggesting different mechanisms of action. Simvastatin treatment resulted in an upregulation of cholesterol synthesis/import genes (including *HMGCR, EBP, DHCR24* and *LSS, LDLR*) and a downregulation of cholesterol export genes (*ABCA1/G1*) (Fig. 3e). This aligns with the activation of the SREBP2-mediated cholesterol homeostasis feedback-loop, employed to minimise cholesterol depletion, following HMG-CoA reductase inhibition [49], in line with this we observed only a moderate reduction in cholesterol levels of 9-18% (Fig. 3g). Simvastatin treatment also led to the upregulation of marker genes associated with active ALK2/BMP signalling (e.g. ID1/3 and SMAD5), further supporting a novel role for ALK2 in cholesterol homeostasis (Fig. 3e). By contrast, ALK2 inhibitor treatment leads to the complete downregulation of cholesterol synthesis genes (Fig. 3e), including the transcription factor *SREBF2,* which encodes SREBP2, indicating an impaired feedback-loop which leads to a greater reduction of intracellular cholesterol of 11-48% (Fig. 3g).We observe a concomitant increase in *NR1H3 and SREBF1* and expression which encode the transcription factors LXR-alpha and SREBP1, respectively, another LXR target gene associated with fatty acid and lipid metabolism [50]. We also observed an increase in *MYLIP* expression (Fig. 3e), which encodes IDOL an E3 ubiquitin ligase inducible degrader of low-density lipoprotein receptor (LDL-R) (Fig. 3f) [51], and a subsequent decrease in LDL-R protein expression (Extended data Fig.3g).

Following combinatorial treatment of M4K2009 or TP-0184 plus simvastatin the suppression of BMP signalling expression is maintained, however there are stark differences in the expression of cholesterol homeostasis genes and activation of *SREBF2* (Fig. 3e). Combinatorial treatment involving M4K2009 more effectively supresses the feedback loop activated by simvastatin, limiting the re-expression of cholesterol biosynthesis genes, and maintains increased cholesterol export (*ABCA1*) and decreased import (*MYLIP*) (Fig. 3f). By contrast, the changes in expression following the TP-0184 combination treatment more closely resemble the changes observed following simvastatin treatment alone. In line with this, we see the largest reduction of intracellular cholesterol levels following M4K2009/statin combinatorial treatment (70-80%) in three *ACVR1-*mutant DMG models (Fig. 3g and Extended data Fig.3h).

Given the potent synergy observed *in vitro* we next assessed the efficacy of M4K2009 plus simvastatin *in vivo* using an orthotopically implanted *ACVR1*-mutant patient-derived model (HSJD_DIPG_007-AKAluc) (Fig. 3h). Initially, *in vivo* tolerability performed in NSG mice determined the maximum tolerated dose (MTD) of both M4K2009 and simvastatin to be 50 mg/kg (Extended data Fig.3i). PK analysis showed that 20 mg/kg and 50 mg/kg simvastatin resulted in comparable concentrations in the brain pons (∼10 μM, 10-fold greater than *in vitro* GI_50_ determinations ∼0.1–1 μM)(Extended data Fig.3j), we therefore decided to proceed with the lower dose to minimise risk of toxicity with prolonged treatment. Single-agent treatment significantly extended the median survival by 10% compared to vehicle; simvastatin p=0.0073 and M4K2009 p=0.0003 (Fig. 3i). The combinatorial treatment arm significantly extended the median survival by 59 days (20%) compared to vehicle (356 days vs 297, p=0.0031), by 20 days compared to M4K2009 which trended towards significance (356 vs 336 days, p=0.0695), and by 17 days compared to simvastatin (356 vs 339 days, p=0.9215), and significantly lowered BLI compared to vehicle (p=0.0254) (Fig. 3i-j). Whilst providing encouraging support for the use of combined M4K2009 and statins in *ACVR1*-mutant DMG, all mice eventually succumb to the disease, highlighting the need to more fully understand the underlying mechanism of response.

### ALK2 inhibitor effects on DMG cells can be rescued with exogenous cholesterol

We hypothesised that the dampened response to combination therapy *in vivo*, compared to the strong synergy scores *in vitro*, may be due to a microenvironmental supply of cholesterol by normal astrocytes. We therefore performed concentration-response curves for each inhibitor with or without exogenous cholesterol supplemented in the media, in both human (*in vitro*) and mouse (*ex vivo*) models. This resulted in a dramatic dose-dependent rescue of cell viability following only M4K2009 (∼1-2 log-fold increase in GI_50_), but not TP-0184 or statin treatment (Fig. 4a-b). In addition, the strong synergistic interaction we observed between ALK2 inhibitors and statins was lost in a dose-dependent manner with increasing concentrations of exogenous cholesterol, most notably in the context of the M4K2009 combinations (Fig. 4c-d). To assess the contribution of cholesterol synthesis versus export as modulators of this phenotype we measured both intra- and extracellular (detected in media) cholesterol and caspase 3/7 activation (apoptosis marker) over time. M4K2009 plus simvastatin combinatorial treatment resulted in the greatest reduction of intracellular cholesterol levels and concomitant increase in extracellular cholesterol over time, which was not seen for TP-0184 (Extended data Fig.4a-b), peaking at 96h and coinciding with caspase 3/7 activation (Extended data Fig.4c-d), with single-agent effects higher in HSJD_DIPG_007 compared to SU_DIPG_IV.

**Figure 4:**
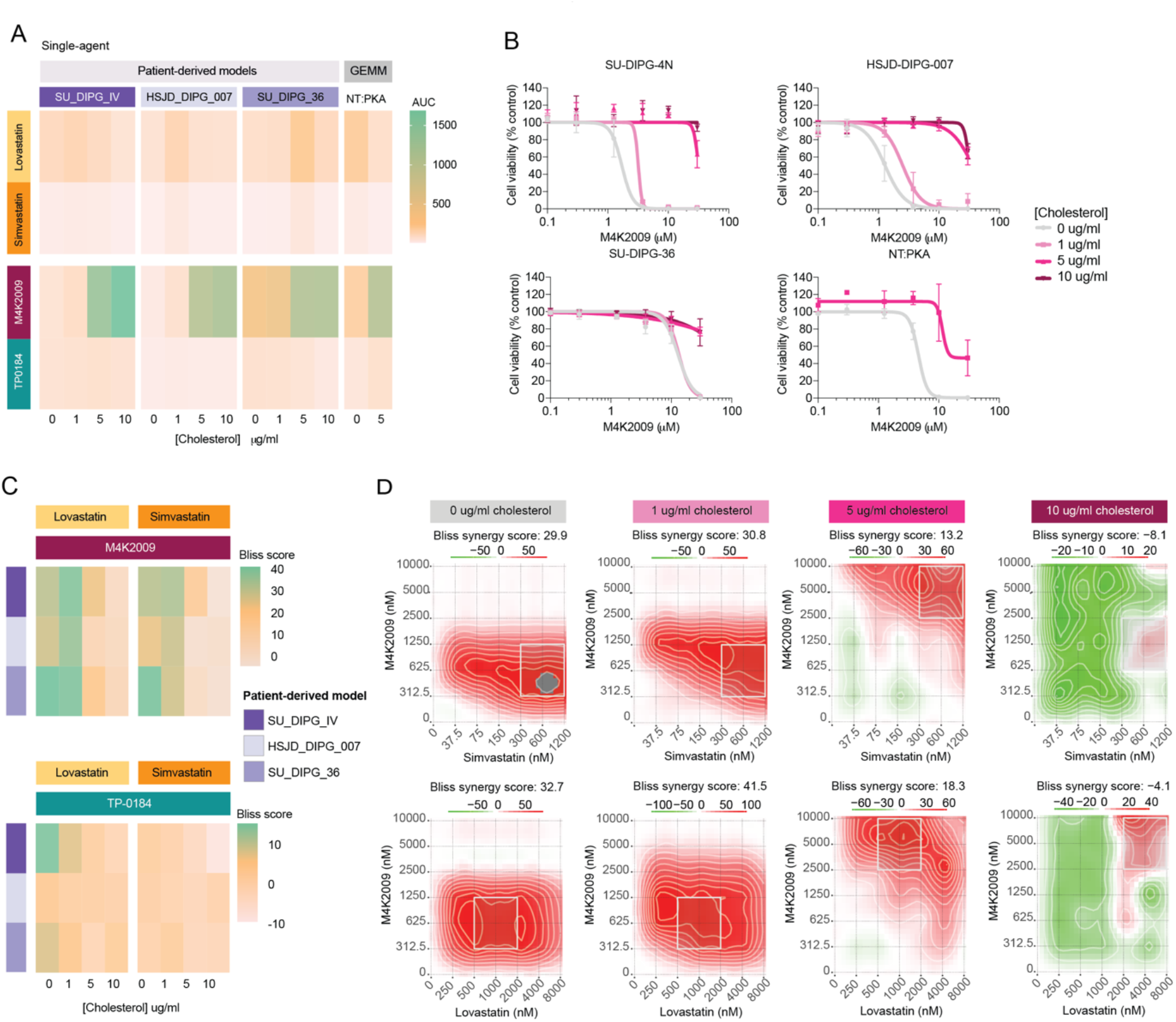
ALK2 inhibitor effects on DMG cells can be rescued with exogenous cholesterol. **A,** Heatmap of mean AUC calculated from concentration-response curves for lovastatin, simvastatin, M4K2009 and TP-0184 with cells seeded in normal SCM or with cholesterol supplemented SCM (at 1, 5 or 10 μg/ml) in three patient-derived *ACVR1*-mutant models and a *ACVR1*-mutant GEMM model (GEMM n=2, patient-derived modes n=3). The AUC is plotted by colour according to the key provided. **B,** Concentration-response curves for M4K2009 with cells seeded in normal SCM or with cholesterol supplemented SCM (at 1, 5 or 10 μg/ml) in three patient-derived *ACVR1*-mutant models and a *ACVR1*-mutant GEMM. Data shown as mean ± SD (GEMM n=2, patient-derived modes n=3). Curves were fitted using [inhibitor] vs response, variable slope, four parameters in Prism. **C,** Heatmap of Bliss scores for M4K2009 or TP-0184 plus lovastatin (left) or simvastatin (right) with cells seeded in SCM or in SCM supplemented with cholesterol (at 1, 5 or 10 μg/ml) in three patient-derived *ACVR1*-mutant models. (n=3). **D,** Representative Bliss synergy plots showing M4K2009 (y-axes) in combination with simvastatin (x-axes, top) or lovastatin (x-axes, bottom) with increasing concentrations of cholesterol supplemented SCM (from left to right). Plots generated using Synergy Finder 3.0. Bliss scores > 10 (red) indicate synergy and <10 (green) indicate antagonism and are plotted according to the key provided.

### Differentiation of cells to an AC-like cell state abrogates the sensitivity to ALK2 inhibition

We next determined if shifting cell-state proportions from OPC-like towards an AC-like population impacts the response to single-agent or combination treatment with ALK2 inhibitor and/or statins by supplementing stem-cell media (SCM) or base media with 10% FBS to drive differentiation *in vitro* [10, 13]. HSJD_DIPG_007 cells were cultured for > 20 passages in FBS-containing media and then characterised with respect to their morphology, growth kinetics, cell-type markers (immunofluorescence) and gene expression (scRNAseq) (Fig. 5a). Cells cultured with FBS showed clear morphological changes towards larger, flatter cells (Fig. 5b-c) with slower growth-kinetics (Fig. 5d) and a significant increase in basal intracellular cholesterol (Fig. 5e). We also observed a significant increase in markers associated with the astrocytic (GFAP/vimentin [52–54]) and neural (MS1[55]) phenotypes and decrease in those associated with oligodendrocytes and stemness (OLIG2/PDGFRA/SOX2 [10, 56, 57]), compared to parental (Fig. 5f and Extended data Fig.5a).

**Figure 5:**
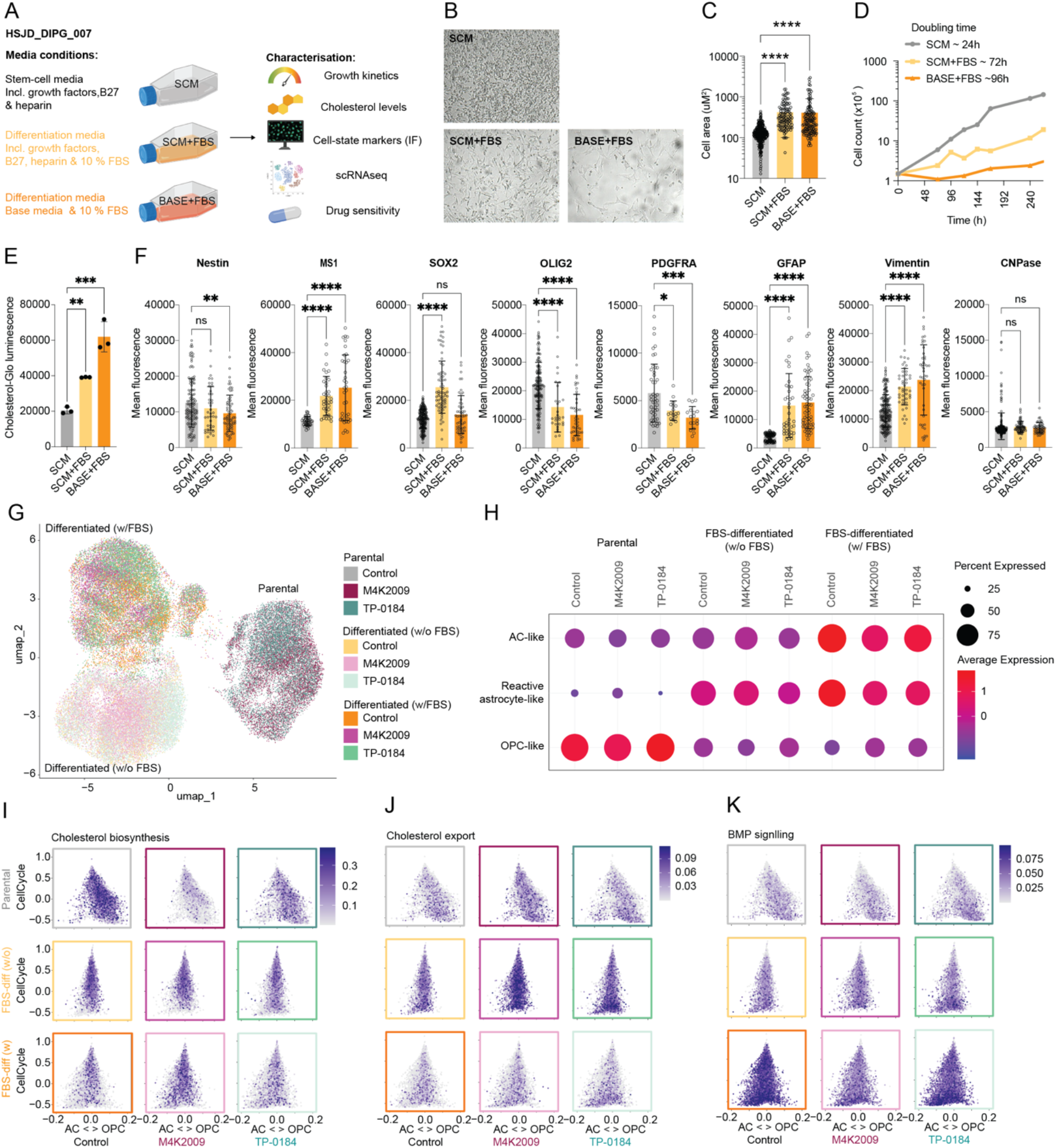
Differentiation of cells to an AC-like cell state abrogates the sensitivity to ALK2 inhibition. **A,** Outline of the different FBS-differentiation media conditions and the subsequent downstream characterisation performed. **B,** Brightfield images (10x objective) of HSJD_DIPG_007 cells cultured in the indicated media. **C,** Quantification of individual cell area (cm^2^) of HSJD_DIPG_007 cells cultured under different media conditions using QuPath. p values calculated by one-way ANOVA (**** p<0.0001) compared to SCM condition. Dots represent individual cells. **D,** Growth curves and calculated doubling times for HSJD_DIPG_007 cells cultured in the indicated media. **E,** Cholesterol-Glo luminescence measuring intracellular cholesterol levels in HSJD_DIPG_007 cells cultured under different media conditions. Cells were seeded in three technical replicates at 10,000 cells in duplicate 96-well plates for 16 hours. Data shown as mean ± SD (n=3). p values calculated by unpaired t-test (**p<0.01, *** p<0.001) compared to SCM condition. **F,** Quantification of the mean immunofluorescence staining of cell markers (nestin, MS1, SOX2, OLIG2, PDGFRA, GFAP, vimentin and CNPase) in HSJD_DIPG_007 cells cultured under different media conditions. p values calculated by one-way ANOVA (* p<0.05, ** p<0.01, *** p<0.001 **** p<0.0001) compared to SCM condition. Dots represent individual cells. **G,** UMAP representation of scRNA-seq data for HSJD_DIPG_007 cells cultured under different media conditions; FBS-differentiated cells were seeded for this experiment both with FBS (differentiated w/ FBS) and without FBS (differentiated w/o FBS) and treated with GI_50_ concentrations of drug (or vehicle DMSO) for 24 h. **H,** Bubble plot showing the AC-like, reactive astrocyte-like and OPC-like expression signature scores for each culture condition with or without drug treatment. The average signature expression is plotted by colour according to the key provided. Circle size indicates the percentage of cells expressing the gene signature. **I,** 2D-representation of the cell cycle scores (y-axis) and astrocyte (AC-like) vs. oligodendrocyte progenitor cell (OPC-like) scores (x-axis) for all single nuclei, coloured by their cholesterol biosynthesis expression score for each culture condition and drug treatment. **J,** 2D-representation of the cell cycle scores (y-axis) and astrocyte (AC-like) vs. oligodendrocyte progenitor cell (OPC-like) scores (x-axis) for all single nuclei, coloured by their cholesterol export expression score for each culture condition and drug treatment. **K,** 2D-representation of the cell cycle scores (y-axis) and astrocyte (AC-like) vs. oligodendrocyte progenitor cell (OPC-like) scores (x-axis) for all single nuclei, coloured by their BMP signalling expression score for each culture condition and drug treatment.

Single-cell RNAseq analysis was performed on parental (SCM) and FBS-differentiated (SCM+FBS only) HSJD_DIPG_007 cells to further assess cell-state changes and response to acute ALK2 inhibitor treatment (24h). The differentiated cells were seeded both in the presence or absence of cholesterol containing FBS to determine if extracellular cholesterol modulates expression changes alone or following ALK2 inhibitor treatment. Analysis revealed the culture condition as the primary driver of sample separation within the UMAP projection (Fig. 5g) and differences in cell-state proportions (Fig. 5h and Extended data Fig.5b). In the FBS-differentiated cultures we observed an increase in the proportion of cells expressing AC-like and reactive astrocyte-like signatures [58] and a decrease in cells expressing an OPC-like signature compared to parental, irrespective of ALK2 inhibitor treatment (Fig. 5h and Extended data Fig.5b). Interestingly, in the FBS-differentiated cells seeded with FBS (i.e. high extracellular cholesterol) short-term treatment with ALK2 inhibitors leads to a moderate reduction in the proportion of AC-like and reactive astrocyte-like cells and increase in OPC-like cells compared to control (Fig. 5h). 2D tri-plot representations highlight the shift in cell-state from a more OPC-like population in the parental cells to more AC-like in the differentiated cells which is associated with altered cholesterol and BMP-signalling gene expression signatures (Fig. 5i-k). ALK2 inhibitor treatment decreases cholesterol biosynthesis in the more OPC-like parental cells (Fig. 5i, top row) but not in the more AC-like FBS-differentiated cells (Fig. 5i, middle and bottom row) compared to their respective control. Additionally, FBS-differentiated cells show a reduction in cholesterol synthesis compared to parental (Fig. 5i, left column), except in those treated with M4K2009 where we observe an increase (Fig. 5i, middle column). We observed a unique increase in cholesterol export in the FBS-differentiated cells, seeded without FBS, most pronounced following M4K2009 treatment (Fig. 5j, middle row). Interestingly, the culture condition alone was sufficient to alter BMP-signalling (Fig. 5k). The FBS-differentiated cultures seeded with FBS showed increased expression of BMP-signalling genes, including the canonical downstream effectors ID1/ID3 and the ALK2-mutant ligand activin A (Extended data Fig.5c), which was only partially reduced by M4K2009 treatment, suggesting that the greater extracellular availability of cholesterol activates BMP signalling in DMG cells potentially serving as a feedback loop (Fig. 5k, bottom row). Additionally, the ALK2 wild-type ligand BMP7 was upregulated in cells seeded without FBS (Extended data Fig.5c), suggesting an alternative mechanism might be at play in AC-like cells when extracellular cholesterol is scarce.

This shift in cell-state proportions coincided with a decrease in the single-agent M4K2009 but not TP-0184 or simvastatin sensitivity (Extended data Fig.5d-f) and a decrease in the Bliss synergy score for M4K2009 plus simvastatin (Extended data Fig.5g), as assessed in FBS-free SCM media. Additionally, normal human astrocyte cultures derived from both frontal lobe (HA-FL) or the brainstem (HA-BS) showed a lack of sensitivity to M4K2009 treatment (GI_50_ >30 μM) but were notably sensitive to TP-0184 treatment (GI_50_ 1-2 μM), similar to DMG cultures (Extended data Fig.5h-i).

### Targeting the microenvironmental supply of cholesterol through modulation of export

Given the rescuing effects of exogenously supplied cholesterol, we next assessed the efficacy of the drug combination treatments using *in vitro* co-cultures and *ex vivo* organotypic brain slices (Fig. 6a). We assessed the interaction between ALK2 inhibitors and simvastatin under different co-culture conditions, using DMG cells (GFP-labelled) grown initially with astrocytes, oligodendrocytes or neural stem cells seeded at a non-proliferative/low density. We observed a significant increase in SU_DIPG_IV GFP integrated intensity only when co-cultured with astrocytes, compared to monoculture, following M4K2009 plus simvastatin treatment at multiple time points (Fig. 6b and Extended data Fig.6a-b). We hypothesised that the provision of cholesterol by astrocytes and subsequent import by the DMG cells might be driving this reduction in drug sensitivity, which may be overcome by promoting its export. To test this, DMG-astrocyte co-cultures were treated with a triple combination of ALK2 inhibitors and simvastatin and the brain-penetrant LXRα-partial/LXRβ-full agonist LXR-623 which promotes cholesterol export via upregulating ABCA1 expression [59]. We observed significant increases in DMG proliferation over the time course (GFP integrated intensity) in the co-culture conditions (compared to DMG mono-culture) treated with M4K2009 plus simvastatin (Fig. 6c, top row), but not TP-0184 plus simvastatin (Fig. 6c, bottom row). The same was observed for M4K2009 and TP-0184 triple combinations (plus simvastatin and LXR623) in HSJD_DIPG_007-GFP, but only at the earlier time points for the M4K2009 triple combination in SU_DIPG_IV-GFP (Fig. 6c). Taken together, LXR agonism, and increased cholesterol export, is able to dampen the protection from the double combination provided by astrocytes.

**Figure 6:**
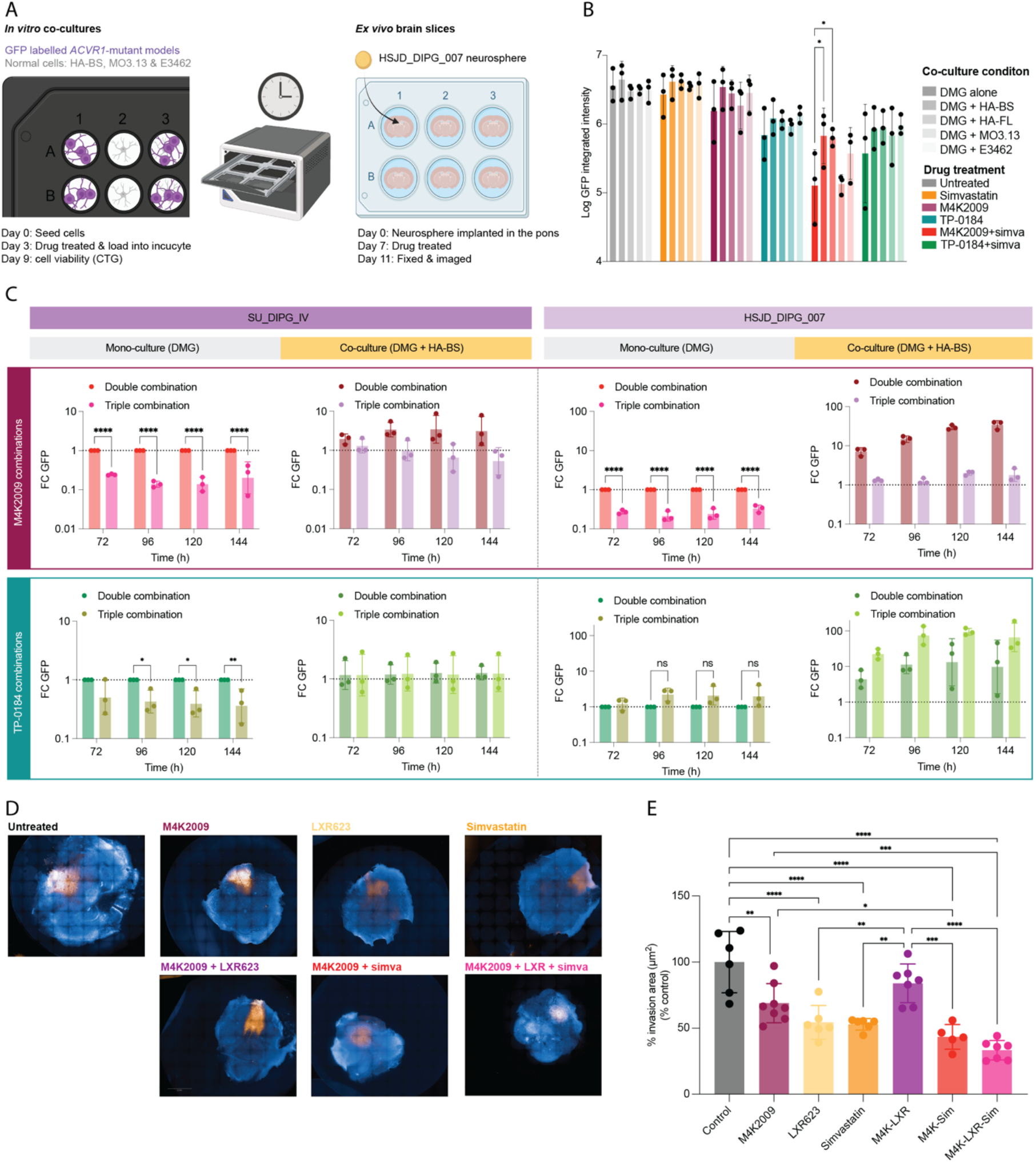
Targeting the microenvironmental supply of cholesterol through modulation of export. **A,** Schematic detailing the conditions of the *in vitro* DMG co-culture assay (using GFP labelled SU_DIPG_IV and HSJD_DIPG_007) and the *ex vivo* organotypic brain slice experiments (using HSJD_DIPG_007 neurospheres). **B,** Log GFP integrated intensity for SU_DIPG_IV-GFP cultured alone or in co-culture following 96 h drug treatment. Data shown as mean ± SD (n=3). p values calculated by two-way ANOVA (* p<0.05). **C,** Fold-change of integrated GFP intensity signal for SU_DIPG_IV-GFP (left) and HSJD_DIPG_007-GFP (right) compared to mono-culture following M4K2009 (top) or TP-0184 (bottom) combination treatment. Data shown is mean ± SD (n≥2). p values calculated by one-way ANOVA (* p<0.05, ** p<0.01, **** p<0.0001). **D,** Representative images of coronal slices of normal mouse brain implanted in the pontine region with HSJD_DIPG_007 neurospheres, treated with 0.3 μM M4K2009, 2.5 μM simvastatin, 5 μM LXR623 or double or triple combinations compared with vehicle control (DMSO) for 96 h. Stained with anti-human nuclei antibody (Orange) and counterstained with Hoechst33342 (blue). Scale bar, 2mm. **E,** Quantification of the area invaded by HSJD_DIPG_007 neurospheres calculated using Image J. Data shown is mean ± SD of at least five independent slices. p values calculated by one-way ANOVA (*p>0.05, **p>0.01, *** p<0.001, **** p<0.0001).

Finally, we explored this triple combination in the more physiologically relevant *ex vivo* organotypic brain slices using patient-derived HSJD_DIPG_007 neurospheres implanted into the pons [60]. We observed a significant decrease in the area invaded following treatment with each single-agents (30-50%) and M4K2009 plus simvastatin (60%) compared to vehicle (DMSO) control (Fig. 6d-e). The greatest reduction in invasion was observed following the triple combination treatment (70%), showing significantly lower invasion compared to vehicle, M4K2009 alone, M4K2009 plus LXR623 but not compared to M4K2009 plus simvastatin (Fig. 6e). Thoughtful and thorough optimisation of dosing and scheduling will be required to avoid potential toxicities *in vivo*, still these promising data supports a move towards more complex triple combination approaches.

## Discussion

Targetable epigenetic, metabolomic and genetic dependencies have been identified in DMG-H3K27 and are beginning to enter clinical trials [61–65]. However, despite a decade passing since the discovery that 20-25 % of these tumours harbour somatic *ACVR1* mutations [18–21], which impart a clear dependence on the BMP signalling pathway and sensitivity to multiple chemotypes of ALK2 inhibitors [24, 30, 66], there are still no selective agents available clinically for patients. Although consistent across models and compounds tested [29, 30, 32], preclinical efficacy of these agents is not curative, so alongside the clinical development of ALK2 inhibitors rational drug combination partners are also urgently needed. To address this, we used unbiased multi-omics and multiple drug combination CRISPR knock-out screens to gain deeper understanding of the consequences of ALK2 inhibition in *ACVR1*-mutant patient-derived models and identify novel and mechanistically driven drug combinations.

Together, drug combination CRISPR screens identified novel dependencies in multiple genes encoding key players involved in cholesterol homeostasis. Targeting these hits using small molecule inhibitors of multiple nodes of the cholesterol biosynthesis pathway in combination with ALK2 inhibitors confirmed a synergistic interaction, most notably with the more selective clinical development candidate M4K2009 [24]. Through multi-omic profiling of *ACVR1*-mutant patient-derived DMG models, we identify ALK2 inhibition to lead to downregulation of cholesterol synthesis gene expression via *SREBF2* (encoding SREBP2, a key cholesterol homeostasis transcription factor [35]) and leading to a reduction in cholesterol and increase in the precursory metabolite desmosterol and by-product 3-hydroxy-3-methylglutarate, generated from 3-hydroxy-3-methylglutaryl CoA hydrolysis [48]. Inhibition of HMG-CoA reductase by statins leads to enzymatic reduction of intracellular cholesterol which activates the SREBP2 transcriptional positive feed-back loop to increase cholesterol synthesis gene expression and restore cholesterol levels. This also increases expression of the BMP pathway effectors *ID1/ID2*, suggesting the presence of a previously unrecognised feedback loop via ALK2 in response to low cholesterol which may mediate the high degree of synergy seen with ALK2 inhibition and statins.

The role of BMP signalling and the different BMP ligands is context dependent spanning tumour suppression/promotion, differentiation dynamics and, in atherosclerosis and osteoporosis, lipoprotein/cholesterol metabolism [67–70]. Simvastatin treatment has been shown to increase BMP2, which signals via ALK3/6 [71], serving as a positive cholesterol feedback loop in osteoblasts [69]. The ALK family members ALK5 and ALK1 been associated with regulating cholesterol efflux [72] or mediating LDL uptake in endothelial cells [73], respectively. ALK1 and ALK2 expression has also been linked to high-density lipoprotein levels in aortic endothelial cells [74]. Our data provides evidence towards a previously unappreciated link between ALK2 signalling and the regulation and expression of key cholesterol synthesis and export proteins leading to profound effects on cholesterol homeostasis in DMG.

We also observed transcriptional differences between cells treated with simvastatin plus M4K2009 or TP0814, with the more selective M4K2009 combination more effectively limiting the re-expression of cholesterol biosynthesis genes and cholesterol import (via IDOL regulation of LDL-R) and maintaining cholesterol export (via ABCA1). This transcriptomic response tracked with intracellular cholesterol levels, where M4K2009 plus simvastatin treatment was most effective at reducing cholesterol. Together these data suggest that ALK2 elicits multifactorial effects on transcriptional regulation, export and import of cholesterol such that targeting the receptor renders the cell more vulnerable to enzymatic inhibition of the subsequently lowly expressed cholesterol synthesis proteins; combination of ALK2 inhibitor and statins leads to a critical depletion of intracellular cholesterol that cannot be sufficiently replenished by biosynthesis. *In vivo,* this combination resulted in a significant increase in median survival supporting translation as a novel rational treatment option; while statins are widely used and well-tolerated in clinic [75–78], clinical availability of a selective ALK2 inhibitor is still urgently required.

Cholesterol metabolism has previously been reported as a dependency in both DMG-H3K27 and adult GBM [13, 79–83]. DMG-H3K27 is a heterogenous disease comprised primarily of developmentally stalled OPC-like cells, which are able to self-renew/proliferate, and to a lesser proportion more differentiated astrocytic cell (AC)-like and mesenchymal (MES)-like cells [10]. The cholesterol dependency identified in DMG-H3K27 has recently been attributed to the proliferating OPC-like population of cells [13]. Here, we show that FBS-mediated differentiation from an OPC-like to an AC-like/reactive astrocyte-like (astrocytes responding/adapting to external stimuli/injury [84]) cell-state is associated with decreased cholesterol biosynthesis yet cells maintain high intracellular cholesterol, likely through import of extracellular cholesterol in the FBS, accounting for the significant reduction in sensitivity to selective ALK2 inhibitor single-agent treatment and in combination with statins. This protection via exogenous cholesterol was mirrored in co-culture experiments with normal astrocytes, as previously suggested in the adult GBM setting whereby cholesterol dependency has been linked to increased uptake of cholesterol from the microenvironment, rather than altered biosynthesis, taking advantage of the limitless supply of *de novo* synthesised cholesterol in the brain [82, 85, 86]. Using a similar approach to Villa and colleagues in adult GBM [82], use of the clinical BBB-penetrant LXR agonist LXR623, which promotes cholesterol export [59], was able to mitigate this exogenous cholesterol rescue of the ALK2 inhibitor/statin combination both *in vitro* an *ex vivo*, suggesting the possible utility of a triple drug combination targeting both tumour cell intrinsic and extrinsic mediated genetic/metabolic dependencies.

The present study has several limitations which will be addressed in future work. We have focused much of our assessment of ALK2 inhibitors in *ACVR1*-mutant models, due to their clear dependency on BMP signalling, but we also observe synergistic interactions between ALK2 inhibitors and statins in a wider panel of PDHGG, independent of *ACVR1-*mutation. It will be important to assess BMP signalling dependency in these models as it has been reported to extend beyond *ACVR1* mutation status in DMG-H3K27 [70]. We have also focused our *in vivo* efforts towards assessing the clinically-used statins to target cholesterol synthesis but ongoing efforts are aimed at evaluating the impact of targeting the different nodes of this pathway in order to prioritise the most effective cholesterol targeting agent. Additional experiments to assess the impact of the combination treatment on the immune microenvironment, determine optimal dosing schedules and interactions with radiotherapy and other clinically progressing targeted agents are also warranted [61, 63, 64].

In summary, we identify a novel link between signalling via ALK2 in *ACVR1*-mutant DMG-H3K27 and cholesterol homeostasis, which suggests a rational combinatorial strategy for clinical translation. Statins are attractive agents in this context, given their well-established clinical safety profiles in both adults and children [75–78]. Notably they may also exert a protective effect given the lower incidence of brain tumours and lower associated risk of brain tumours in patients on long-term statin treatment [87–89]. As selective ALK2 inhibitors approach the first early phase clinical trials for patients with *ACVR1*-mutant DMG-H3K27, plans to safely incorporate cholesterol biosynthesis inhibitors into combination studies appears to be a priority.

## Acknowledgements

We acknowledge Brain Tumour Research funding to the Centre of Excellence at ICR. This work was further supported by Children with Cancer UK, Cancer Research UK (DRCRPG-Nov21/100002), the Brain Tumour Charity, Ollie Young Foundation, CRIS Cancer Foundation, Lucas’s Legacy, Doing it for Daniel, Finlay’s Fighters, Alex’s Lemonade Stand Foundation, Cancer Therapeutics Innovation Pipeline program at the Ontario Institute for Cancer Research (OICR), Agora and M4Kpharma and the DIPG Collaborative. The DIPG Collaborative includes The Cure Starts Now Foundation, Brooke Healey Foundation, Melina Michelle Edenfield Foundation, The Cure Starts Now Australia, The Cure Starts Now Canada, Reflections Of Grace Foundation, Yuvaan Tiwari Foundation, Cure Brain Cancer Foundation, Aubreigh’s Army Foundation, Aidan’s Avengers, Run DIPG, Musella Foundation, Love4Lucas Foundation, Whitley’s Wishes, Anna’s Bake Sale Foundation, The Ayla Foundation, The Isabella and Marcus Foundation, Love, Chloe Foundation, Lauren’s Fight for Cure, Robert Connor Dawes Foundation, Ryan’s Hope, The Gold Hope Project, Abby’s Corner Foundation, The DIPG/DMG Collaborative and Snapgrant.com. We acknowledge NHS support for the Biomedical Research Centre at the ICR and Royal Marsden NHS trust. We are grateful for technical support from the ICR Genomics and Proteomics Facilities. We acknowledge Dave Smil, Methvin Isaac and Ahmed Aman for compound supply and kinase-panel data. We acknowledge Louise Howell and Emma Westland for their support with the confocal microscopy analysis. We further acknowledge Fondazione Umberto Veronesi (fellowship to Giulia Pericoli). For the purpose of Open Access, the author has applied a CC BY public copyright licence to any Author Accepted Manuscript (AAM) version arising from this submission.

## Methods

### Primary cell cultures

Patient-derived cultures were established as previously described [90]. Briefly, cells were incubated at 37°C in a humidified atmosphere of 5% CO2. Cultures maintained as 2D were passaged into flasks pre-coated with 2 ng/μl Cultrex laminin (Biotechne, 3446-005-01) while neurospheres were passaged into ultra-low attachment flasks. Accutase solution was used to detach cells from the flask or dissociate neurospheres, cells were incubated with accutase for 2-5 minutes at 37°C. Media was then added at a 1:1 ratio before centrifugation at 1300 rpm for 3 minutes. Cells were re-suspended in medium and split at an appropriate concentration. Cells were routinely STR profiled and checked for mycoplasma.

Patient-derived cultures were grown in complete stem cell media and fed with ∼30% fresh media every 2-3 days. Initially, base media is prepared which is composed of 250 ml DMEM/F12 (Thermo Fisher Scientific, 11330-038), 250 ml Neurobasal-A Medium (Thermo Fisher Scientific, 10888-022), 10 mM HEPES Buffer Solution (Thermo Fisher Scientific, 15630-080), 1 mM MEM Sodium Pyruvate Solution (Thermo Fisher Scientific, 11360-070), 0.1 mM MEM Non-Essential Amino Acids Solution (Thermo Fisher Scientific, 11140-050) and 1x Glutamax-I Supplement (Thermo Fisher Scientific, 35050-061). The complete media is then prepared by supplementing base media with B-27 Supplement Minus Vitamin A 1:50 (Thermo Fisher Scientific, 12587-010), 20 ng/ml recombinant Human-EGF (2B Scientific LTD, Oxford, UK, 100-26), 20 ng/ml recombinant Human-FGF (2B Scientific LTD, 100-146), 10 ng/ml recombinant Human-PDGF-AA (2B Scientific LTD, 100-16), 10 ng/ml recombinant Human-PDGF-BB (2B Scientific LTD, 100-18), and 2 μg/ml Heparin Solution (Stem Cell Technologies, Cambridge, UK, 07980).

### Non-tumour cell cultures

The immortal human-human hybrid cell line MO3.13 that expresses phenotypic characteristics of primary oligodendrocytes was obtained from Cederlane Labs (CLU301) and maintained in high-glucose DMEM (D5796) plus 10% FBS (Gibco, A5670801). Human astrocytes from the cerebral cortex (HA-FL, cat 1800) or brain stem (HA-BS, cat 1840) were obtained from ScienCell. Cells were culture 2 μg/cm^2^ poly-L-lysine (SC-0413 Caltag Medsystems) in defined astrocyte medium (cat 1801, ScienCell). The neural stem cell line E3462-striatum, dissected from a foetal brain of 15 week gestation, was a kind gift from Steve Pollard (University of Edinburgh). E3462 was maintained in DMEM/HAMS-F12 (D8437), 1.5 mg/ml glucose (Sigma G8644), 0.1 mM MEM Non-Essential Amino Acids Solution, 0.012% BSA solution (Gibco 15140-122), 0.1 mM b-mercaptoethanol (Gibco 31350-010), B-27 Supplement Minus Vitamin A 1:100, N2 supplement 1:200 (Gibco 17502-048).

### RCAS:Nestin-Tv-a cell model

The Nestin-Tv-a; Trp53^fl/fl^; Hist1h3b^K27M^, Acvr1^R206H^ (NT:PKA) model has been previously described [32]. Cells were incubated at 37°C in a humidified atmosphere of 5% CO_2_. Cells were maintained as neurospheres and were passaged into ultra-low attachment flasks. Accutase solution was used to dissociate neurospheres for 2-5 minutes at 37°C. Media was then added at a 1:1 ratio before centrifugation at 1300 rpm for 3 minutes. Cells were re-suspended in medium and split at an appropriate concentration. Cells were routinely STR profiled and checked for mycoplasma. Cells were grown in Dulbecco’s Modified Eagle Medium (DMEM) supplemented with 10% proliferation supplement (Stem Cell Technologies), 1 % Pen–Strep (Invitrogen), 20 ng/mL human basic FGF (Invitrogen), 10 ng/mL human EGF (Invitrogen), and 2 μg/mL heparin (Stem Cell Technologies).

### FBS differentiated cell cultures

HSJD_DIPG_007 cells were cultured in complete stem cell media plus 10% FBS (Gibco, A5670801) (SCM+FBS) or base media plus 10% FBS (BM+FBS) for > 10 passages before downstream characterisation including immunofluorescence of cell state markers, growth dynamics, drug sensitivity and transcriptomics.

### Growth curves and optimum seeding density

For growth curves, cells were seeded at 300,000 cells in multiple T25 flasks (pre-coated with laminin for 2D cultures), incubated at 37°C and fed with ∼30 % fresh media every 2-3 days. Every 24 h cells are detached with Accutase as previously described and counted on the Countess III (Thermo Fisher Scientific). Cell number was then plotted on a semi-log graph and doubling times were calculated. For seeding density, cells were seeded at a range of seeding densities in multiple 96- or 384-well plates in 100 μl or 40 μl media, respectively, and incubated at 37°C. Every 24 h cell viability (proportional to cell number) was assessed using CellTiter-Glo (Promega, G7570). Data was analysed in Prism 10.

### Drugs and compounds

M4K2009 was supplied by M4K Pharma. All compounds were purchased from Selleckchem except M4K2163 (Biotechne), LDN-193189/Lovastatin/Simvastatin (Merck/Sigma-Aldrich), TP-0184 (MedChemExpress), Tamoxifen (Cambridge Bioscience), Trametinib (Stratech), LDN-213844 (Medkoo) and Palovarotene (Biomol). See Supplementary Table 7 for details for all compounds and drugs used.

### Kinase-profiling

TP-0184 selectivity was profiled against the Eurofins KinaseProfiler panel (>445 kinases) at 1μM. LDN-193189 selectivity was profiled against the Nanosyn KinomeScan panel at 0.1 and 1 μM [33]. M4K2009 and M4K2163 selectivity were profiled against the Eurofins KinaseProfiler panel (384 kinases) at 1 μM. Data is presented as % inhibition and plotted using CORAL (http://phanstiel-lab.med.unc.edu/CORAL/).

### Concentration-response curves and GI_50_ determinations

Cells were plated at their optimum seeding densities in 100 or 40 μl media in 96- or 384-well plates, respectively, and incubated at 37°C for 72 h. For 96-well assays, test compound was manually added at a range of concentrations (serially diluted 1:2) in 100 μl medium. For 384-well assays, test compound was added at a range of concentrations using the ECHO acoustic liquid handler (Labcyte, Beckman Coulter). Following addition of compound, plates were incubated at 37°C for 72 or 192 h and cell viability was assessed using the CellTiter-Glo assay (Promega). Data were normalised to vehicle (DMSO) control and analysed using Prism 10. GI_50_ values (drug concentration required to reduce the cell number to 50% of vehicle control) were determined using the sigmoidal, 4PL, X is concentration non-linear curve fitting model. AUC was calculated with the baseline set to Y=50.

### Bliss synergy score determinations

Cells were plated at their optimum seeding densities in 40 μl media in 384-well plates and incubated at 37°C for 72 h. Drug combinations were performed as outlined on SynergyFinder 3.0 (https://synergyfinder.fimm.fi), drugs were dispensed using the ECHO acoustic liquid handler. Plates were then incubated at 37°C for 192 h and cell viability was assessed using the CellTiter-Glo assay (Promega). Data were normalised to vehicle control and analysed using the web application SynergyFinder 3.0 using the Bliss synergy model to generate both overall and peak synergy scores [91].

### Drugs and tools library combination screens

The ICR drugs and tools library was designed and constructed in-house and consisted of 869 compounds in total which spanned four 384-well source plates. Compound details and Robust Z-scores are detailed in Supplementary Table 4. Cells were plated at 125 cells/well in 40 μl media in 384-well plates and incubated at 37°C for 72 h. Each compound was dispensed using the ECHO acoustic liquid handler at a single final assay concentration of 1 μM in duplicate plates, one set of these plates were then treated with a sub-lethal concentration of M4K2009 or TP-0184 (final assay concentration of 1 μM). Plates were then incubated at 37°C for 192 h and cell viability was assessed using the CellTiter-Glo assay. Data was normalised to vehicle DMSO control wells, the difference in cell viability (% control) was calculated for each compound with or without M4K2009/TP-0184 treatment. Data is mean ± SD for three biological replicates.

### Genome-wide CRISPR/Cas9 screens at the Institute of Cancer research

Lentivirus production: the human GeCKOv2 CRISPR knockout pooled library (123411 sgRNAs), kindly provided by Dr. Feng Zhang (Addgene #1000000048) [92], was transfected with lipofectamine 3000 into HEK293FT (RRID:CVCL_6911) cells along with psPAX2 (Addgene #12260, RRID:Addgene_12260) and pMD2.G plasmids (Addgene #12259, RRID:Addgene_12259) to produce lentivirus. Approximately 5 h after transfection, media was replaced with fresh media. Lentivirus supernatant was harvested 72 h post-transfection and filtered using 0.45 µM steriflip filter unit (Merck). Lentivirus was concentrated using Lenti-X concentrator solution (Clontech) and stored at -80 °C.

GeCKOv2 CRISPR-Cas9 genome-wide screens: genome-wide screens were performed using a minimum of 8.6×10^5^ cells seeded onto laminin coated T175 flasks. Cells were transduced at a MOI of <0.3 with the human GeCKOv2 library (123,411 sgRNAs[92]), in the presence of polybrene 8 μg/ml, for a minimum of 500-fold coverage. Media was refreshed 16 h post-transduction and cells were incubated for a further 48 h. Media was then replaced with fresh media containing 5 μg/ml puromycin and cells were incubated for 48 h to allow for efficient gene-editing. At this point (5 days post-transduction) T0 samples were collected and stored at -80°C for later processing. The remaining cells were cultured for a further 10 doublings (23 days) at 500-fold coverage in the absence of antibiotics, fresh media with or without ALK2i (0.3μM LDN193189, 0.4μM M4K2009). was added to cells every 2-3 days and cells were passaged once a week. T10 samples (10 doublings) were collected and stored at -80°C for genomic DNA extraction.

Fastq files were analysed with MAGeCK (0.5.9.5) [93] and mapped to the GeCKOv2 guide reference library using the count function with trim-5 enabled. Maximum-likelihood estimation (MLE) testing was carried out on raw read counts with default parameters using the mle function. Z-score values were obtained and a difference in Z-scores between control and treated experiment arms was calculated by subtracting the control Z-scores from respective treated scores.

### Pathway-level agreement analysis of CRISPR screens

Gene-level Z-scores from three CRISPR screens were filtered to include genes common across all datasets. Gene set enrichment analysis was performed using Hallmark pathways from msigdbr and the fgsea package, ranking genes by −1*Z score. Normalized enrichment scores (NES) were calculated for each pathway and experiment, and Pearson correlation of pathway NES profiles was used to quantify similarity between screens. Results were visualized as a correlation heatmap.

### Genome-wide CRISPR/enCas12a screens at the Hudson Institute of Medical Research

The SU_DIPG_IV and SU_DIPG_36 cell lines were transduced with enCas12 (pRDA_174, Addgene #136476) and selected with blasticidin (5ug/ml) to select for Cas12 expression. Cas12 activity was validated as a 50% reduction in GFP fluorescence 7 days after transduction with a lentiviral construct co-expressing GFP and a gRNA targeting GFP, pRDA_221 (Addgene #169142). Cells were infected using Humagne Set C, D genome-wide Cas12 libraries (Broad Institute, Addgene #172650, #172651) at MOI<0.3, selected with puromycin 2 ug/ml for one week before addition of ALK2 inhibitor (TP-0184). Screening was conducted at 750x representation in the presence and absence of 0.4μM TP-0184 for 2-3 weeks. Genomic DNA isolation, amplicon processing and NGS was conducted as previously described [94].

### Panel CRISPR/Cas9 screens at the Montreal Neurological Institute/McGill University

SU_DIPG_21, SU_DIPG_36, and HSJD_DIPG_007 cell lines were engineered to stably express Cas9 by infection with Lenti-Cas9-blast (Addgene #52962), followed by blasticidin selection. Robust Cas9 activity was confirmed by transducing cells with pKLV2-U6gRNA5(gGFP)-PGKmCherry2AGFP-W (Addgene #67982) or pKLV2-U6gRNA5(Empty)-PGKmCherry2AGFP-W (Addgene #67981), followed by flow cytometry to measure the expression of mCherry and GFP (which is blunted in cells expressing Cas9 and sgGFP). The CRISPR screens were performed using a custom library encoding 13,243 sgRNA targeting 3319 genes that encode “druggable” molecules, in addition to controls [95]. For each cell line, the transduction conditions were optimized to achieve a multiplicity of infection (MOI) of 0.3-0.4, and pilot screens were performed to identify a concentration of M4K2163 that decreases growth or viability by approximately 20% (IC_20_) for the duration of the screen. The following M4K2163 concentrations were selected: SU_DIPG_21: 0.2μM; SU_DIPG_36: 0.2μM; HSJD_DIPG_007: 0.1μM. For each cell line, 2 x 10^8^ cells were transduced in suspension at the optimized MOI in the presence of 17 μg/mL polybrene. The next day, the infection medium was replaced with selection medium containing 1 μg/mL puromycin. Three days later, the cells were harvested, and 8×10^6^ cells were distributed in four 15 cm plates, in three replicates. After six additional days of culture to allow the depletion of sgRNAs encoding essential genes, each replicate was split in two. M4K2163 was added at the optimized concentrations to one half, and an equivalent volume of vehicle (DMSO) to the other half. The cells were passaged and exposed to fresh drug or vehicle every 4 days thereafter. At each time point, 7 x 10^6^ cells were passaged to maintain >500X coverage for each sgRNA encoded in the library. The screens were continued until approximately 15-18 cell doublings in the vehicle condition (SU_DIPG_21: 39 days; SU_DIPG_36: 26 days; HSJD_DIPG_007: 22 days). The genomic DNA was extracted with the QIAamp DNA Mini Kit (Qiagen), RNAse-treated, and quantified. Library preparation and sequencing was performed at the Princess Margaret Genomics Centre (Toronto, Canada). The reads were mapped to sgRNA sequences encoded in the library, quantified, and normalized to the total read counts. sgRNA enrichment and depletion between drug- and vehicle-treated cells was assessed using DrugZ [96].

### Gene knock-out nucleofection

Each gene was targeted using two independent gRNA and HDR templates (Integrated DNA Technologies. Coralville, IA) (Supplementary Table 8). crRNA and tracrRNA (200 μM each) were co-incubated to form guide RNA (gRNA) complexes (95°C for 5 minutes) according to the manufacturer’s instructions. gRNA complexes were incubated with 20 μM recombinant Cas9 protein (Integrated DNA Technologies) at 37°C for 15 minutes to assemble ribonucleoprotein (RNP) complexes. RNP complexes (3 μl) were electroporated together with corresponding HDR templates (1 μl, 200 μM) into 2×10^5^ SU_DIPG_IV, SU_DIPG_36 or HSJD_DIPG_007 cells in 20 μl of SE “X” nucleofection buffer using a 4D-Nucleofector X kit (Lonza, Cat#V4XP-3032) and unit 16-well nucleocuvette (Lonza Group LTD, Basel, Switzerland) with program CA-137. After nucleofection, cells were equilibrated in pre-warmed rescue media and plated into 6-well (3 ml) and 96-well plates (100 μl). Cells plated into 6-well plates were expanded for downstream analysis. Cells plated into 96-well plates were incubated for 72 h at 37°C then treated with a range of concentrations of ALK2 inhibitor (in 100 μl) and incubated at 37°C for 192 h. Cell viability was assessed using CellTiter-Glo assay (Promega). Single-gene knockout efficiency was validated on genomic DNA using Tracking of Indels by Decomposition (TIDE) analysis.

### CRISPR knockout efficiency assessment using TIDE

To assess gene editing efficiency, genomic DNA was extracted from electroporated cells and genomic regions of interest were amplified by polymerase chain reaction (PCR). PCR products were purified using AMPure XP beads as per the manufacturers instructions (Beckman Coulter, A63881). DNA was sequenced at DNA sequencing & services, University of Dundee, Scotland. The efficiency of Cas9-mediated gene editing was determined by comparing DNA sequences from target gene-specific gRNA-nucleofected cells versus unedited controls (no RNP, Cas9 alone, and *AAVS1*) using the TIDE analysis tool [97].

### Nucleic acid extraction

DNA and RNA were extracted following the DNeasy Blood & Tissue kit (QIAGEN, Hilden, Germany, 69504) and the RNeasy Plus Mini Kit protocols (QIAGEN, 74134), respectively. Occasionally a dual RNA/DNA extraction kit, Quick-DNA/RNA™ Miniprep Plus Kit (Zymo Research, Irvine CA, USA, D7003T), was also used following manufacturer’s instructions. Concentrations were measured using the Qubit dsDNA Assay Kits (Thermo Fisher Scientific, Q32850, Q32851) and/or a TapeStation 4200 (Agilent, Santa Clara CA, USA).

### Whole-exome sequencing

Libraries were prepared from 1000ng of DNA using the NEBNext® Ultra™ II DNA Library Prep Kit according to the manufacturer’s instructions (New England Biolabs, E7103). The resultant libraries were captured with xGen Exome Hyb Panel V2 (Integrated DNA Technologies, 10005151) following the IDT hybridization capture protocol. Samples were sequenced on an Illumina NovaSeq X Plus system using the NovaSeq™ X Series 25B 300 cycle Reagent Kit with a 150bp PE sequencing to 100x or 200x raw coverage (Illumina, 20104706).

Capture reads were aligned to the GRCh37 build of the human genome using bwa bwa v0.7.12 (bio-bwa.sourceforge.net). The Genome Analysis Tool Kit (GATK, v4 .1.9.0) and PicardTools (v2.23.8, <u>picard.sourceforge.net</u>), were used for PCR duplicate removal, base quality score recalibration and generation of metrics for each sample. Single nucleotide variants (SNVs) and small insertions/deletions (indels) were called using current GATK best practices and somatic mutations were identified by joint calling with Mutect2 (broadinstitute.org/gatk/). Variants were annotated using the Ensembl Variant Effect Predictor v101 (ensembl.org/info/docs/variation/vep). Copy number was obtained by calculating log_2_ ratios of tumour/normal coverage binned into exons of known Ensembl genes, smoothed using circular binary segmentation (DNAcopy, <u>v1.84.0</u>), and processed using in-house scripts in R.

### RNA-sequencing

RNA sequencing was performed by Source Genomic Services (Cambridge, UK). mRNA libraries were prepared from 500ng of RNA using the NEBNext® Ultra™ II RNA Library Prep Kit (New England Biolabs, E7770L) with the NEBNext® Poly(A) mRNA Magnetic Isolation Module (New England Biolabs, E7490L), according to the manufacturer’s instructions. Samples were sequenced on an Illumina NovaSeq X system using the NovaSeq™ X Series 25B 300 cycle Reagent Kit with 150bp PE sequencing to either 30 million or 120 million read pairs per sample (Illumina, 20104706).

RNA-seq data were aligned with STAR (v 2.7.11a) and read counts within ensembl genes collected using HTSeq (v 2.0.3). Following rlog transformation and normalization, differential expression was assigned with DESeq2 (v1.50.2). GSEA was carried out using the R package fgsea (v1.36.0) based on curated canonical pathways (MsigDB, Broad Institute)

### Methylation profiling

DNA methylation array was performed at the University College London Genomics Centre (London, UK). A total of 500ng DNA was bisulphite-modified and analysed for genome-wide methylation patterns using the Illumina Human MethylationEPIC 850k beadarray, according to the manufacturer’s instructions (Illumina).

Methylation data from the Illumina Infinium HumanMethylation850 BeadChip were preprocessed using the minfi package in R (v1.56.0). The Heidelberg brain tumor classifier (MNP12.8; molecularneuropathology.org [36]) was used to assign a calibrated score to each case, associating it with one of the 185 tumor entities that feature within the current classifier. Clustering of β values from methylation arrays was performed based on correlation distance using a Ward algorithm. DNA copy number was derived from combined log_2_ intensity data based on an internal median processed using the R packages minfi (v1.56.0) and conumee (v1.44.0) to call copy number in 15,431 bins across the genome.

### Bulk transcriptomic analysis

Cells were seeded in T75 flasks at 4×10^5^ – 1×10^6^ cells/flask, to achieve ∼70% confluence at end-point, and incubated at 37°C for 72 h. Cells were then treated with GI_50_ concentrations of single-agent compound for 24 h. The GI_50_ values were determined using a 72 h concentration-response assay; M4K2009 was added at a final assay concentration of 8 μM for all models and TP-0184 was added at 8, 2 and 1.5 μM for SU_DIPG_IV, SU_DIPG_36 and HSJD_DIPG_007, respectively. For assessing combinatorial drug treatment response using bulk RNA sequencing SU_DIPG_IV cells were treated with either 8 μM M4K2009/TP-0184, 0.5 μM simvastatin or a combination of 8μM ALK2 inhibitor plus 0.5 μM simvastatin for 24 h. Following the 24 h drug treatment, cells were scraped from the flasks in their media (+/- drug) and centrifuged at 1300 rpm for 3 minutes. Cell pellets were washed once in ice-cold PBS, centrifuged at 1300 rpm for 3 minutes and snap frozen in dry ice. Three biological replicates were sent for bulk RNA sequencing analysis. RNA extraction was performed as outlined above. Accession number

### Single-cell RNAseq

HSJD_DIPG_007 cells cultured in stem cell media (SCM) or SCM plus FBS were seeded in T75 flasks at 0.5 -2×10^6^ cells/flask and incubated at 37°C for 72 - 192 h until ∼ 60% confluent. Cells were treated with 8 μM M4K2009 or 1 μM TP-0184 for 24 h. For FBS cultures, cells were treated with and without FBS in the media (removed 24 h prior to treatment). Each sample was prepared as a single cell suspension, cells were gently detached with Accutase (#A6964, Sigma), washed and resuspended with complete media, counted and then washed again in PBS with 0.04% BSA. Single cell suspensions were processed through the 10X Chromium Single Cell Platform using Chromium Single Cell 3’ v3.1 Chemistry (10X Genomics, Pleasanton, CA) following the manufacturer’s protocol. A total of 5000 cells were added to each channel of a chip to be partitioned into Gel Beads in Emulsion (GEMs) in the Chromium instrument, followed by cell lysis and barcoded reverse transcription of RNA in the droplets. Breaking of the emulsion was followed by amplification, fragmentation, and addition of adaptor and sample index. Single Cells 3’ Gene Expression libraries were sequenced on the NovaSeq 6000/Novaseq X Plus and processed with Cell Ranger analysis pipeline.

### Proteomics

Cells were seeded in T75 flasks at 4×10^5^–1×10^6^ cells/flask, to achieve ∼70% confluence at end-point, and incubated at 37°C for 72 h. Cells were then treated with GI_50_ concentrations of single-agent compound for 24 h. The GI_50_ values were determined using a 72 h concentration-response assay; M4K2009 was added at a final assay concentration of 8 μM for all models and TP-0184 was added at 8, 2 and 1.5 μM for SU_DIPG_IV, SU_DIPG_36 and HSJD_DIPG_007, respectively. Following the 24 h drug treatment, cells were scraped from the flasks in their media (+/- drug) and centrifuged at 1300 rpm for 3 minutes. Cell pellets were washed once in ice-cold PBS, centrifuged at 1300 rpm for 3 minutes and snap frozen in dry ice. Three biological replicates were sent for proteomic analysis.

Cell pellets were lysed in a buffer containing 1% sodium deoxycholate (SDC), 100 mM triethylammonium bicarbonate (TEAB), 10% isopropanol and 50 mM NaCl, freshly supplemented with 5 mM TCEP (Thermo, Bond-breaker), 10 mM iodoacetamide, universal nuclease 1:2000 vol/vol (Pierce, #88700) and Halt protease and phosphatase inhibitor cocktail (Thermo, #78442, 100X) with 5 min of bath sonication. Protein concentration was measured with the Quick Start Bradford protein assay (Bio-Rad). Aliquots of 30 μg of total protein were digested overnight with trypsin (Pierce, 1:20) at room temperature. Peptides were labelled with the TMTpro reagents (Thermo) by adding 5 μL of the reagent (25 μg/μL) into 12.5 μL of sample volume. The TMTpro mixture was acidified with formic acid at 2% and the precipitated SDC was removed by centrifugation.

The peptide pool was fractionated with high pH Reversed-Phase chromatography using the XBridge C18 column (2.1 × 150 mm, 3.5 μm, Waters) on an UltiMate 3000 HPLC system over a 1% gradient in 35 min. Mobile phase A was 0.1% (v/v) ammonium hydroxide and mobile phase B was 0.1% ammonium hydroxide (v/v) in acetonitrile.

The LC-MS analysis of the 12plex set was performed on an UltiMate 3000 system coupled to the Orbitrap Lumos mass spectrometer (Thermo). Peptides were loaded onto the Acclaim PepMap 100, 100 μm × 2 cm C18, 5 μm, trapping column at a flow rate of 10 μL/min and analysed with an Acclaim PepMap (75 μm × 50 cm, 2 μm, 100 Å) C18 capillary column connected to a stainless-steel emitter (Thermo, ES542). Mobile phase A was 0.1% formic acid and mobile phase B was 80% acetonitrile, 0.1% formic acid. The separation method was as follows: for 90 min gradient 5%-38% B, for 10 min up to 95% B, for 5 min isocratic at 95% B, re-equilibration to 5% B in 5 min, for 10 min isocratic at 5% B at a flow rate of 300 nL/min. MS scans were acquired in the range of 375-1,500 m/z with a mass resolution of 120,000. Precursors were selected in the top speed mode in 3 sec cycles and isolated for HCD fragmentation with quadrupole isolation window 0.7 Th. Collision energy was 36% with AGC 1×10^5^ and maximum injection time 86 ms at 50,000 resolution. Targeted precursors were dynamically excluded from further fragmentation for 45 sec with 7 ppm mass tolerance.

The LC-MS analysis for the 18plex set was performed on a Vanquish Neo HPLC system (Thermo) coupled to the Orbitrap Ascend mass spectrometer (Thermo) using a 25 cm capillary column (Waters, nanoE MZ PST BEH130 C18, 1.7 μm, 75 μm × 250 mm) over a 110 min gradient 5%-35% of mobile phase B composed of 80% acetonitrile, 0.1% formic acid. Peptides were preconcentrated onto a PepMap 100, C18, 5 μm, 0.3×5 mm, 1500 bar, trapping column following elution in the analytical column which was attached to a Nanospray Flex ion source via a stainless-steel emitter. MS spectra were collected at Orbitrap mass resolution of 120,000 and precursors were selected for HCD fragmentation in the top speed mode (3 sec) with collision energy 32% and iontrap detection in turbo scan rate. MS3 scans were triggered by Real Time Search (RTS) against a fasta file containing UniProt Homo sapiens reviewed canonical and isoform sequences with Synchronous Precursor Selection (SPS 10 notches) and HCD fragmentation with collision energy 65% at 45,000 Orbitrap resolution. Targeted precursors were dynamically excluded from further activation for 45 sec with 10 ppm mass tolerance and RTS close-out was enabled with max 4 peptides per protein. Static modifications for RTS were TMTpro16plex at K/n-term (+304.2071), Carbamidomethyl at C (+57.0215) and variable modifications were Deamidated NQ (+0.984) and Oxidation of M (+15.9949) with maximum 1 missed-cleavage and 2 variable modifications per peptide.

The 12plex raw data were processed with the Sequest HT node in Proteome Discoverer 2.4 (Thermo) and the 18plex data were processed with the Sequest HT and Comet nodes in Proteome Discoverer 3.0 for protein identification and quantification against a fasta file containing reviewed UniProt Homo sapiens entries. The precursor mass tolerance was set at 20 ppm and the fragment ion mass tolerance at 0.5 Da (or 1 Da for Comet) with up to 2 trypsin missed-cleavages allowed. TMTpro at N-terminus/K and Carbamidomethyl at C were defined as static modifications. Dynamic modifications were oxidation of M and deamidation of N/Q. Peptide confidence was estimated with the Percolator node and peptide FDR was set at 0.01 based on target-decoy search. Only unique peptides were used for quantification, considering protein groups for peptide uniqueness. Peptides with average reporter signal-to-noise greater than 3 were used for protein quantification.

### Metabolomics

Cells were seeded in T175 flasks at 9.2×10^5^–2.3×10^6^ cells/flask, to achieve ∼70% confluence at end-point, and incubated at 37°C for 72 h. Cells were then treated with GI_50_ concentrations of single-agent compound for 24 h. The GI_50_ values were determined using a 72 h concentration-response assay; M4K2009 was added at a final assay concentration of 8 μM for all models and TP-0184 was added at 8, 2 and 1.5 μM for SU_DIPG_IV, SU_DIPG_36 and HSJD_DIPG_007, respectively. Following the 24 h drug treatment, cells were scraped from the flasks in their media (+/- drug) and centrifuged at 1300 rpm for 3 minutes. Cell pellets were washed once in ice-cold PBS, centrifuged at 1300 rpm for 3 minutes and snap frozen in dry ice. Five biological replicates were sent for metabolomic analysis. Samples were shipped to and processed by Metabolon using their Global Discovery Panel (North Carolina, US).

### Cholesterol rescue assays

Cells were plated at their optimum seeding densities in 40 μl media in 384-well plates and incubated at 37°C for 72 h. Compounds were added at a range of concentrations using the ECHO acoustic liquid handler (Labcyte, Beckman Coulter) in four replicates. Cholesterol (C4951, Sigma/Merck) was then added at 0, 1, 5 or 10 ug/ml (in 10 μl media) to one of the four replicates. Plates were incubated at 37°C for 192 h and cell viability was assessed using the CellTiter-Glo assay (Promega). Data were normalised to vehicle (DMSO) control and analysed using Prism 10. AUC was calculated with the baseline set to Y=50.

### Intra- and extracellular cholesterol determinations

Cholesterol was measured using the Promega Cholesterol-Glo kit (#J3191) according to the protocol. Briefly, cells were plated at their optimum seeding densities in 100 μl in 96-well plates and incubated at 37°C for 72 h. Cells were then treated with a GI_50_ concentration of compound (100 μl) and incubated at 37°C for 24, 48 or 96 h. SU_DIPG_IV and HJSD_DIPG_007 cells were treated with 0.3 μM simvastatin, 3 μM M4K2009 and 3 or 1 μM TP-0184, respectively. Extracellular cholesterol levels were determined using the media (25 μl aliquot/well) while intracellular cholesterol levels were determined following cell lysis. Duplicate plates were used to assess cell viability using the CellTiter-Glo assay. Cholesterol-Glo luminescence data was normalised to

CellTiter-Glo luminescence, and the log fold-change was calculated for each time-point compared to untreated (vehicle DMSO) control.

### DMG co-culture assays

DMG-GFP labelled cells were plated at their optimum seeding densities along with unlabelled non-tumour cells (astrocytes, oligodendrocytes and neural stem cells), at minimally proliferative seeding densities, in 100 μl compete media in 96-well plates and incubated at 37°C for 72 h. Cells were then treated with either single-agent or combinations of 0.3 μM simvastatin, 3 μM M4K2009, 3 μM TP-0184, 4 μM LXR623 in 100 μl complete media. Plates were then placed into a SX5 Incucyte (Sartorius) and incubated at 37°C for subsequent time course analysis. GFP integrated intensity and confluence was exported from the Incucyte analysis software. Duplicate plates were used to allow assessment at end-point of cell viability using the CellTiter-Glo assay and cholesterol levels using the Cholesterol-Glo assay. The fold-change of GFP integrated intensity was calculated for mono-culture vs co-culture for each treatment condition over time. Additionally, time course data was plotted in Prism 10 and the AUC (baseline Y=50) was calculated and normalised to the mono-culture for each treatment condition or all to the untreated mono-culture.

### ABCA1 immunofluorescence

HSJD_DIPG_007, SU_DIPG_IV and SU_DIPG_036 cells were seeded in 96-well plates (Phenoplate, Revvity) at their optimal seeding densities and incubated at 37°C for 72 h. Cells were then treated with GI_50_ concentrations of M4K2009 (8 μM) or TP-0184 (HSJD_DIPG_007, 1.5 μM; SU_DIPG_36, 2 μM; SU_DIPG_IV, 8 μM) or 100 μg/ml desmosterol (Merck/Sigma, D6513) and incubated at 37°C for 48 h. Cells were then fixed with 4% paraformaldehyde at room temperature for 10 minutes and washed three times with phosphate buffered saline (PBS) solution. Cells were permeabilised with 0.5% Triton X-100 solution for 10 minutes at room temperature and then blocked with appropriate serum according to the species of secondary antibody for 1 h at room temperature. Primary antibody directed against ABCA1 (ab18180 clone AB.H10, Abcam, 1:200) was added and incubated overnight at 4°C. Cells were then washed in PBS three times and incubated with Alexa Fluor488-conjugated secondary antibody (A11001, Invitrogen) 1 h at room temperature. Nuclei were counterstained with DAPI and the samples were imaged using a Opera Phenix Plus High Content Screening System and results were analysed using the Harmony software version 5.2. Signal intensity was calculated for each cell.

### Cell marker immunofluorescence

HSJD-DIPG007 cells cultured in stem-cell media with or without FBS were seeded in 8 well-chamber slides (Cole Palmer) pre-coated with laminin in their respective media and incubated at 37°C. Once ∼ 70% confluent, cells were fixed with 4% paraformaldehyde at room temperature for 10 minutes and washed three times with phosphate buffered saline (PBS) solution. Cells were permeabilised with 0.5% Triton X-100 solution for 10 minutes at room temperature and then blocked with appropriate serum according to the species of secondary antibody for 1 h at room temperature. Primary antibodies directed against nestin (MAB5326 clone 10C2, Millipore, 1:400), SOX2 (3579 clone D6D9, Cell Signaling, 1:400), GFAP (Z334, Dako, 1:50), OLIG2 (Ab9610, Millipore, 1:200), Musashi-1/MS1 (Ab5977, Millipore, 1:200), Vimentin (M0725, Dako, 1:100) and PDGFRA (3174 clone D1E1E, Cell Signaling, 1:100) were added and incubated overnight at 4°C. Cells were then washed in PBS three times and incubated with Alexa Fluor488/555-conjugated secondary antibodies (A11001, A11008 & A31572, Invitrogen) 1 h at room temperature. Nuclei were counterstained with DAPI and the samples were imaged using a Zeiss LSM700 confocal microscope. Staining images were quantified using QuPath imaging software version 5.

### Organotypic whole-brain slices (OBS)

All animal procedures were under the European Communities Council Directive N. 2010/63/EU and the Italian Ministry of Health guidelines (DL 26/2014) and approved by the Italian Ministry of Health and by the local Institutional Animal Care and Use Committee at Istituto Superiore di Sanità (Rome, Italy; protocol n. A69A0.N.RMV, 2019). Coronal whole-brain organotypic slices encompassing the pons were prepared from CD1 mice pups (postnatal days 6–7; Charles River) as previously described [39], with some modifications. In brief, mice were decapitated and brains rapidly dissected and placed in ice-cold artificial cerebrospinal fluid containing (in mmol/L): 126 NaCl, 3.5 KCl, 1.2 NaH_2_PO_4_, 1.2 MgCl_2_, 2 CaCl_2_, 25 NaHCO_3_, and 11 glucose (pH 7.3), saturated with 95% O_2_ and 5% CO_2_. The brain was then embedded in 3% SeaPlaque agarose (Lonza) in PBS and 300 μm-thick coronal slices were cut on a vibrating microtome (Campden Instruments), constantly cooled, and oxygenated. Each slice was transferred onto a porous membrane (0.45 μm pore size; Millipore), placed on a Millipore culture insert, and inserted into 6-well plates with 1.2 ml of cell culture medium/well, where the inserts were placed. The slices were incubated at 35°C, 5% CO_2_ for 7 days before the experiments to allow the inflammatory reaction following the mechanical procedure to subside. Following the first day of culture, the medium was replaced with fresh medium and, from that time, changed every 48 h. Seven days after slices were sectioned, HSJD_DIPG_007 parental neurospheres with a diameter of 250 to 300 μm (1 neurosphere/slice) were implanted on the pontine area, and following 3 days of coculture, preparations were treated for 4 days with 0.3 μM M4K2009, 2.5 μM simvastatin, 5 μM LXR623 or double or triple combinations compared with vehicle control (DMSO). After this time, slices were fixed with 10% buffered formalin for 2 h at room temperature (RT) and washed twice with PBS for 10 minutes. Slices were then permeabilized with 1% Triton in PBS for 90 minutes and then blocked with 10% goat serum + 1% BSA + 0.1% Triton for 60 minutes. Slices were incubated overnight with anti-human nuclei antibody (MAB4383, Millipore. 1:300). Slices were washed with PBS twice for 10 minutes and incubated with the Alexa Fluor-555 goat anti-mouse secondary antibody overnight (A21424, Invitrogen. 1:500). Hoechst33342 was used as a counterstain (1:10,000 in PBS for 45 minutes at RT; Invitrogen). Images were taken on an Operetta CLS (PerkinElmer) in confocal mode (10×; z-stack 45 μm). Quantification was performed manually with Image J version 1.34.

### Tolerability and pharmacokinetics

All *in vivo* experiments were approved by the local Animal Welfare and Ethics Review Board at the Institute of Cancer Research and carried out in accordance with the UK Home Office Animals (Scientific Procedures) Act of 1986, the UK National Cancer Research Institute guidelines for the welfare of animals in cancer research, and the ARRIVE (Animal Research: Reporting *In Vivo* Experiments) guidelines. NOD.Cg-*Prkdc^scid^ Il2rg^tm1Wjl^*/SzJ (NSG) were treated with an oral dose (PO) of M4K2009 (50 mg/kg) and Simvastatin (20, 50 and 100 mg/Kg) for 2 or 3 weeks (5 days on, 2 days off). Once maximum tolerated dose was determined for each drug, animals were treated PO with a combination of: M4K2009 (50 mg/Kg) and Simvastatin (20 mg/Kg). Animals were monitored daily. Plasma and brain tissue samples were taken at 2h post-last dose. Analysis was carried out by liquid chromatography-tandem mass spectrometry (LC-MS/MS) using a Waters Xevo TQ-XS coupled with an Acquity UPLC H-class system (Waters, Herts, UK). Chromatography was carried out using a Phenomenex (Macclesfield, UK) Kinetex C18 column (1.7 µm, 50 mm × 2.1 mm). Data acquisition was performed using Targetlynx, version 4.2.

### AKALuc transduction of in vivo model

Neurospheres were dissociated into a single-cell suspension and pre-incubated with 2 μg/ml polybrene for 1 – 2 h before transduction with AKA-luciferase reporter (Venus-labelled, Addgene #124701) lentivirus. Following a 16 h incubation at 37°C media was replaced and cells were incubated for a further 96 h at 37°C. Transduced cells were then chemically selected for using 1 mg/ml Geneticin^TM^ (G418 sulfate) (Gibco 10131035) for 12 days (refreshed every 2–3 days) before the top 10% of Venus-expressing cells were sorted using the FACSymphony S6 cell sorter (BD Biosciences).

### In vivo efficacy studies

A single cell suspension was prepared from HSJD_DIPG_007-Akaluc cells immediately prior to implantation in NSG mice. Animals received pre-operative analgesia consisting of subcutaneous buprenorphine (0.015 mg/kg) and meloxicam (0.5 mg/kg), along with a local subcutaneous injection of bupivacaine/lidocaine along the incision site. Animals were anesthetized with 4% isoflurane and maintained at 2% to 3% isoflurane delivered in oxygen (1 L/min). Mice were placed on a stereotactic apparatus for orthotopic implantation, with coordinates x = +1.0, z = −0.8, y = −4 mm from the lambda used for delivery to the pons. Then, 2-4 μL of cell suspension (250,000 cells) was stereotactically implanted per animal, using a 25-gauge SGE standard fixed needle syringe (SGE 005000) at a rate of rate of 2 μL/min using a digital pump (HA1100, Pico Plus Elite; Harvard Apparatus). At 24 hours post-surgery, animals were administered an additional dose of meloxicam (0.5 mg/kg). HSJD_DIPG_007-Akaluc animals were randomized into four groups based on bioluminescence intensity as determined by IVIS imaging (IVIS Spectrum CT from PerkinElmer): group 1 vehicle, group 2 M4K2009 (50mg/kg), group 3 Simvastatin (20 mg/kg) and group 4 M4K2009 (50mg/kg) and Simvastatin (20mg/kg). Each drug was prepared individually and combined immediately prior to dosing. Drug combinations were administered as a single dose (0.1 mL per 10 g body weight). The vehicles used were as follows: 0.5% methylcellulose (15 cP) in water for M4K2009, and 5% DMSO, 5% Tween-80, 40% of 0.5% methylcellulose (15 cP) in water, and 50% water for Simvastatin. Vehicle-treated control animals received a single dose of the combined vehicle formulation. Animals were treated for up to 20 weeks, 5 days on, 2 days off. Mice were weighed twice a week and sacrificed by cervical dislocation upon deterioration of condition and tissue was taken for further analysis.

## Data availability

Data are available within the article and its Supplementary data files. Bulk RNA sequencing data from patient-derived models (+/- inhibitor treatment) have been submitted to GEO and are accessible under GSE325979. Single-cell RNA sequencing data from parental HSJD_DIPG_007 and FBS-differentiated cultures have been submitted to GEO and are accessible under GSE326085. The mass spectrometry proteomics data have been deposited to the ProteomeXchange Consortium via the PRIDE partner repository with the dataset identifier PXD076157.

## Supplementary information

Supplementary Table 1. Kinase-profile data for ALK2 inhibitors.

Supplementary Table 2. Difference in Z-scores (+/-ALK2i) for the drug combination CRISPR screens.

Supplementary Table 3. Overlap of top 500 hits from each CRISPR screen.

Supplementary Table 4. Difference in robust Z-scores (+/- ALK2i) for the high-throughput drug combination screens in SU_DIPG_IV.

Supplementary Table 5. Summary of genetic variants observed in the panel of paediatric-type diffuse high grade glioma models.

Supplementary Table 6. Metabolomic data for SU_DIPG_IV, SU_DIPG_36 and HSJD_DIPG_007 treated with ALK2 inhibitors.

Supplementary Table 7. Details for compounds and drugs. Supplementary Table 8. crRNA and HDR template sequences.

Extended data Figures 1-6 inserted below.

## Extended data Figures 1-6

**Extended data Fig.1:**
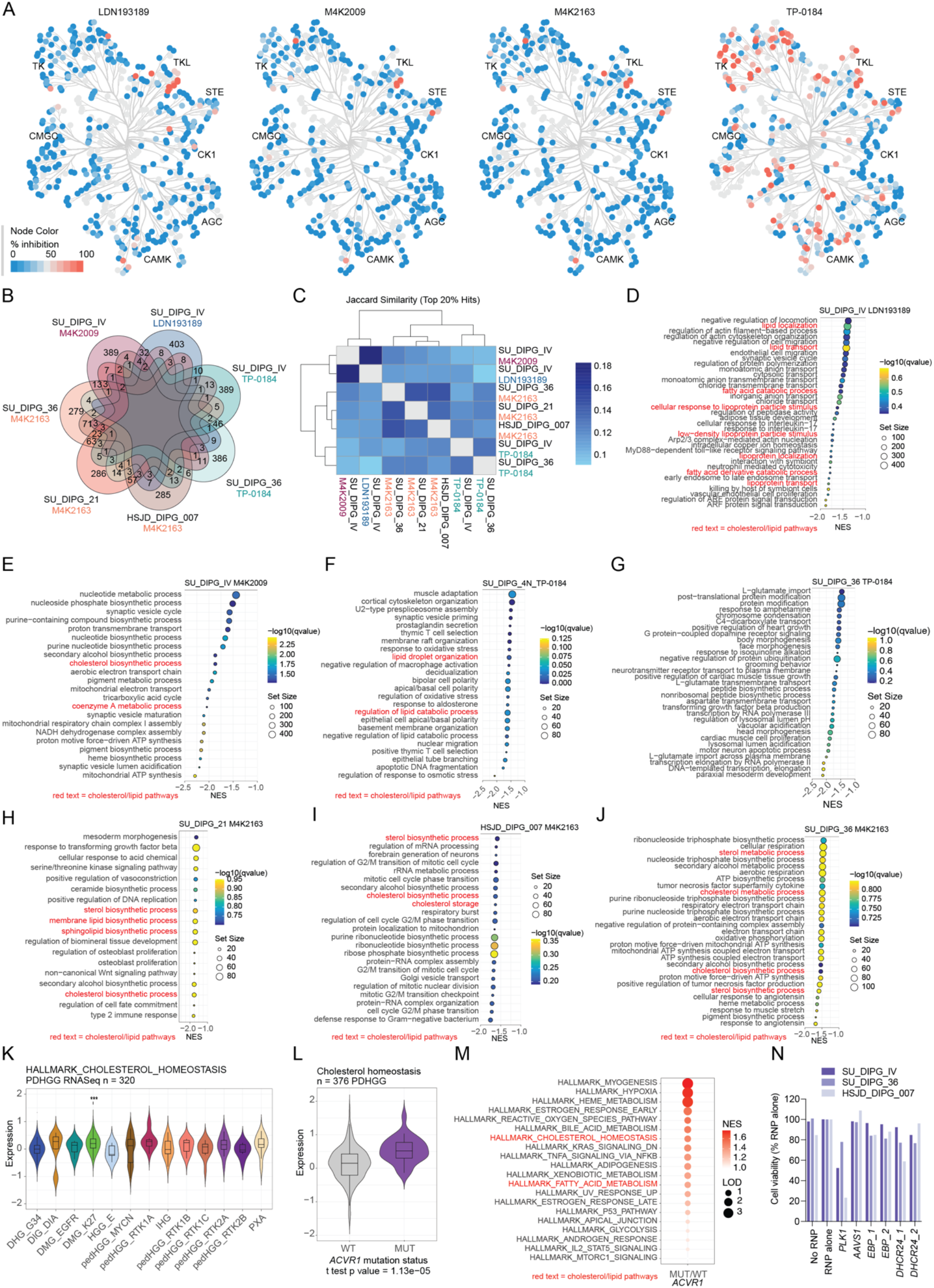
ALK2 inhibitor kinase-profiles, CRISPR screen overlap and gene-set enrichment analysis and bulk tumour expression analysis. **A,** Kinase activity plots for LDN193189, M4K2009, M4K2163 and TP-0184. M4K2009, M4K2163 and TP-0184 were screened across the Eurofins kinase panel and tested at a fixed concentration 1 μM. LDN-193189 selectivity was profiled against the Nanosyn KinomeScan panel at 0.1 μM. Colour and size of node indicate percentage kinase activity inhibition (plotted using CORAL). **B,** Seven-way Venn diagram showing the overlap of ALK2 inhibitor sensitising hits across all CRISPR screens. **C,** Heatmap of the Jaccard similarity score calculated using the top 20 % of ALK2 inhibitor sensitising hits from each CRISPR screen. The Jaccard similarity score is plotted by colour according to the key provided. **D,** Bubble plot of depleted pathways identified by GSEA of the ranked ALK2 inhibitor sensitising hits from the SU_DIPG_IV +/- LDN193189 CRISPR screen. Red text indicates pathways associated with cholesterol/lipid homeostasis. The -log10 (q value) is plotted by colour according to the key provided and the gene set size is indicated by circle size. **E,** Bubble plot of depleted pathways identified by GSEA of the ranked ALK2 inhibitor sensitising hits from the SU_DIPG_IV +/- M4K2009 CRISPR screen. Red text indicates pathways associated with cholesterol/lipid homeostasis. The -log10 (q value) is plotted by colour according to the key provided and the gene set size is indicated by circle size. **F,** Bubble plot of depleted pathways identified by GSEA of the ranked ALK2 inhibitor sensitising hits from the SU_DIPG_IV +/- TP-0184 CRISPR screen. Red text indicates pathways associated with cholesterol/lipid homeostasis. The -log10 (q value) is plotted by colour according to the key provided and the gene set size is indicated by circle size. **G,** Bubble plot of depleted pathways identified by GSEA of the ranked ALK2 inhibitor sensitising hits from the SU_DIPG_36 +/- TP-0184 CRISPR screen. Red text indicates pathways associated with cholesterol/lipid homeostasis. The -log10 (q value) is plotted by colour according to the key provided and the gene set size is indicated by circle size. **H,** Bubble plot of depleted pathways identified by GSEA of the ranked ALK2 inhibitor sensitising hits from the SU_DIPG_21 +/- M4K2163 CRISPR screen. Red text indicates pathways associated with cholesterol/lipid homeostasis. The -log10 (q value) is plotted by colour according to the key provided and the gene set size is indicated by circle size. **I,** Bubble plot of depleted pathways identified by GSEA of the ranked ALK2 inhibitor sensitising hits from the HSJD_DIPG_007 +/- M4K2163 CRISPR screen. Red text indicates pathways associated with cholesterol/lipid homeostasis. The -log10 (q value) is plotted by colour according to the key provided and the gene set size is indicated by circle size. **J,** Bubble plot of depleted pathways identified by GSEA of the ranked ALK2 inhibitor sensitising hits from the SU_DIPG_36 +/- M4K2163 CRISPR screen. Red text indicates pathways associated with cholesterol/lipid homeostasis. The -log10 (q value) is plotted by colour according to the key provided and the gene set size is indicated by circle size. **K,** Violin plot showing the “Hallmark cholesterol homeostasis” bulk-RNA signature expression across 320 PDHGG tumours sub-grouped by methylation classification (MNP12.8). p values calculated using t-test, *** = p<0.001. **L,** Violin plot showing the bulk-RNA expression of the leadingEdge enrichments in “Hallmark cholesterol homeostasis” and “Reactome cholesterol biosynthesis” across 376 DMG-H3K27 tumours split according to *ACVR1*-mutation status. p value calculated using t-test. **M,** GSEA of differentially expressed genes between *ACVR1*-mutant and *ACVR1*-wt patient-derived models. Red text indicates pathways associated with cholesterol/lipid homeostasis. The NES score is plotted by colour according to the key provided and the LOD is indicated by circle size. **N,** Cell viability (y-axis) following CRISPR-Cas9 directed hit gene knockout normalised to RNP alone condition in three *ACVR1*-mutant patient derived models (n=1). *AAVS1* negative gRNA control and *PLK1* positive gRNA control were used.

**Extended data Fig.2:**
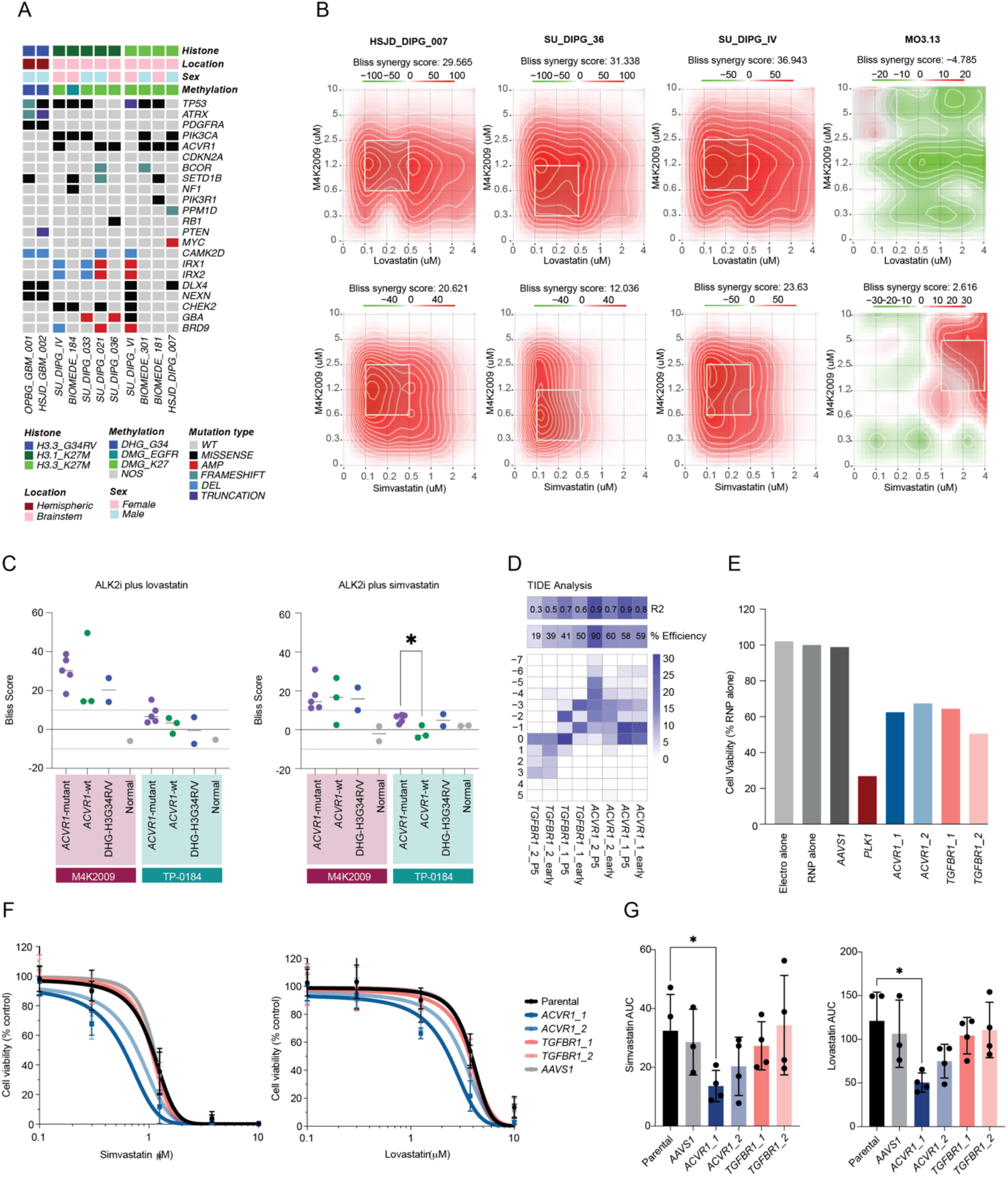
Drug screen validation and *ACVR1/TGFBR1* knock-out models. **A,** Oncoprint representation of the panel of PDHGG patient-derived models used for single-agent and combination drug assays. Samples are arranged in columns with genes labelled along rows. Clinicopathologic and molecular annotations are annotated according to the key provided. **B,** Representative Bliss synergy plots showing M4K2009 (y-axes) in combination with lovastatin (x-axes, top row) or simvastatin (x-axes, bottom row). Plots generated using Synergy Finder 3.0. Bliss scores > 10 (red) indicate synergy and <10 (green) indicate antagonism and are plotted according to the key provided. **C,** Bliss synergy scores for M4K2009 (M4K) or TP-0184 (TP) in combination with lovastatin (left) or simvastatin (right) in a panel of patient-derived PDHGG and non-PDHGG models (MO3.13 and HA-FL) split by subgroup / genotype; *ACVR1*-mutant DMG-H3K27 (purple), *ACVR1*-wt DMG-H3K27 (green), DHG-H3G34R/V (blue) and normal (grey) (n=3). Bliss scores > 10 indicate synergy and <10 indicate antagonism. p values calculated by one-way ANOVA (* p<0.05) compared to *ACVR1*-mutant DMG-H3K27 models. **D,** Heatmap summarising the TIDE analysis of editing efficiency (R2 and % efficiency) of gRNAs targeting *ACVR1* and *TGFBR1* in SU_DIPG_IV cells immediately after nucleofection (early) and after five passages (P5). Frequency of insertions and deletions in base pair positions are scaled in blue. **E,** SU_DIPG_IV cell viability (y-axis) following CRISPR-Cas9 directed *ACVR1* (blue) or *TGFBR1* (pink) knockdown normalised to RNP alone condition (n=1). compared to *AAVS1* negative gRNA control and *PLK1* positive gRNA control were used. **F,** Concentration-response curves for simvastatin (left) and lovastatin (right) in *ACVR1* or *TGFBR1* knockdown SU_DIPG_IV cells (n=2). Data normalised to DMSO vehicle control, shown as mean ± SD. Curves were fitted using [inhibitor] vs response, variable slope, four parameters in Prism. **G,** AUC (y-axis) was calculated from simvastatin (left) and lovastatin (right) concentration-response curves (normalised to DMSO vehicle control, n=2) using Prism (baseline Y=50). p values calculated by unpaired Student’s t test, *p<0.05.

**Extended data Fig.3:**
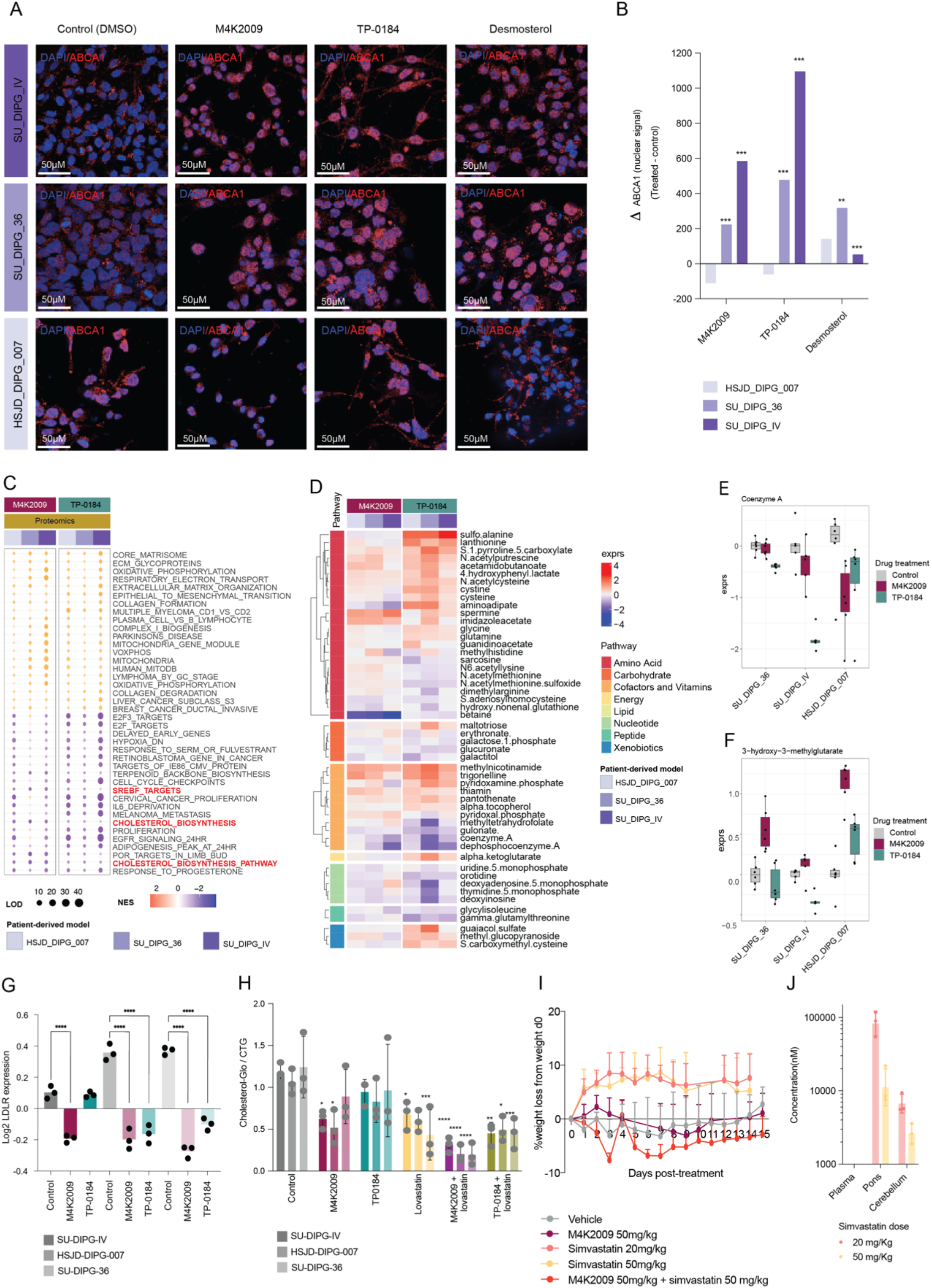
ABCA1 immunofluorescence, multi-omic analysis following ALK2i treatment and *in vivo* tolerability/pharmacokinetics. **A,** ABCA1 immunofluorescence staining for three patient-derived *ACVR1*-mutant models following treatment with vehicle (DMSO) or GI_50_ concentrations of M4K2009 / TP-0184 or 100 ug/ml desmosterol for 48 h. Images were acquired using the Opera Phoenix at 63x magnification and are representative of 3 technical replicates. Scale bar = 50μM. **B,** Barplot showing the difference in mean nuclear-localised ABCA1 IF between untreated (DMSO), ALK2 inhibitor or desmosterol treated conditions (three technical replicates). Differences between treated and untreated groups were assessed using permutation testing. The observed difference in means was compared to a null distribution generated by randomly permuting group labels across the pooled observations. **p<0.01, ***p<0.001. **C,** Bubble plot of enriched or depleted pathways identified by GSEA of differentially expressed proteins across three *ACVR1*-mutant patient-derived models. Red text indicates pathways associated with cholesterol/lipid homeostasis. The NES score is plotted by colour according to the key provided and the LOD is indicated by circle size. **D,** Heatmap showing the metabolites expression in three *ACVR1*-mutant patient-derived models following 24 h treatment with equipotent concentrations of M4K2009 or TP-0184. Data normalised to control (DMSO) condition for each patient-derived model. Data is mean of five biological replicates. The expression is plotted by colour according to the key provided and the metabolite pathways are colour coded as indicated. **E,** Box and whisker plot showing the expression of coenzyme A in three *ACVR1*-mutant patient-derived models following 24 h treatment with equipotent concentrations of M4K2009 or TP-0184. Data normalised to control (DMSO) condition for each patient-derived model. Data shown as mean ± SD (n=5). **F,** Box and whisker plot showing the expression of 3-hydroxy-3-methylglutarate in three *ACVR1*-mutant patient-derived models following 24 h treatment with equipotent concentrations of M4K2009 or TP-0184. Data normalised to control (DMSO) condition for each patient-derived model. Data shown as mean ± SD (n=5). **G,** Log2 expression of LDL-R protein in SU_DIPG_IV, HSJD_DIPG_007 and SU_DIPG_36 following 24 h treatment with equipotent concentrations of M4K2009 or TP-0184. Data shown as mean ± SD (n=3) p values calculated by one-way ANOVA (**** p<0.0001) compared to control (DMSO) condition. **H,** Cholesterol-Glo luminescence following 48 h of single-agent or combination treatment using equipotent concentrations in three *ACVR1*-mutant patient-derived models. Data normalised to CTG luminescence and shown as mean ± SD (n=3). p values calculated by two-way ANOVA (* p<0.05, ** p<0.01, *** p<0.001, **** p<0.0001) compared to control (DMSO) condition. **I,** Tolerability in NSG mice exposed to daily oral treatment (PO) with M4K2009 50 mg/kg, simvastatin 20 or 50 mg/kg or M4K2009 50 mg/kg and simvastatin 50 mg/kg over 15 days as assessed by percentage body weight loss relative to day 0. Data shown as mean ± SD (n>3 mice per group). **J,** Brain and plasma concentrations of simvastatin 2h post-last dose at 20 or 50 mg/kg in non-tumour bearing NSG mice. Data shown as mean ± SD for three mice per condition.

**Extended data Fig.4:**
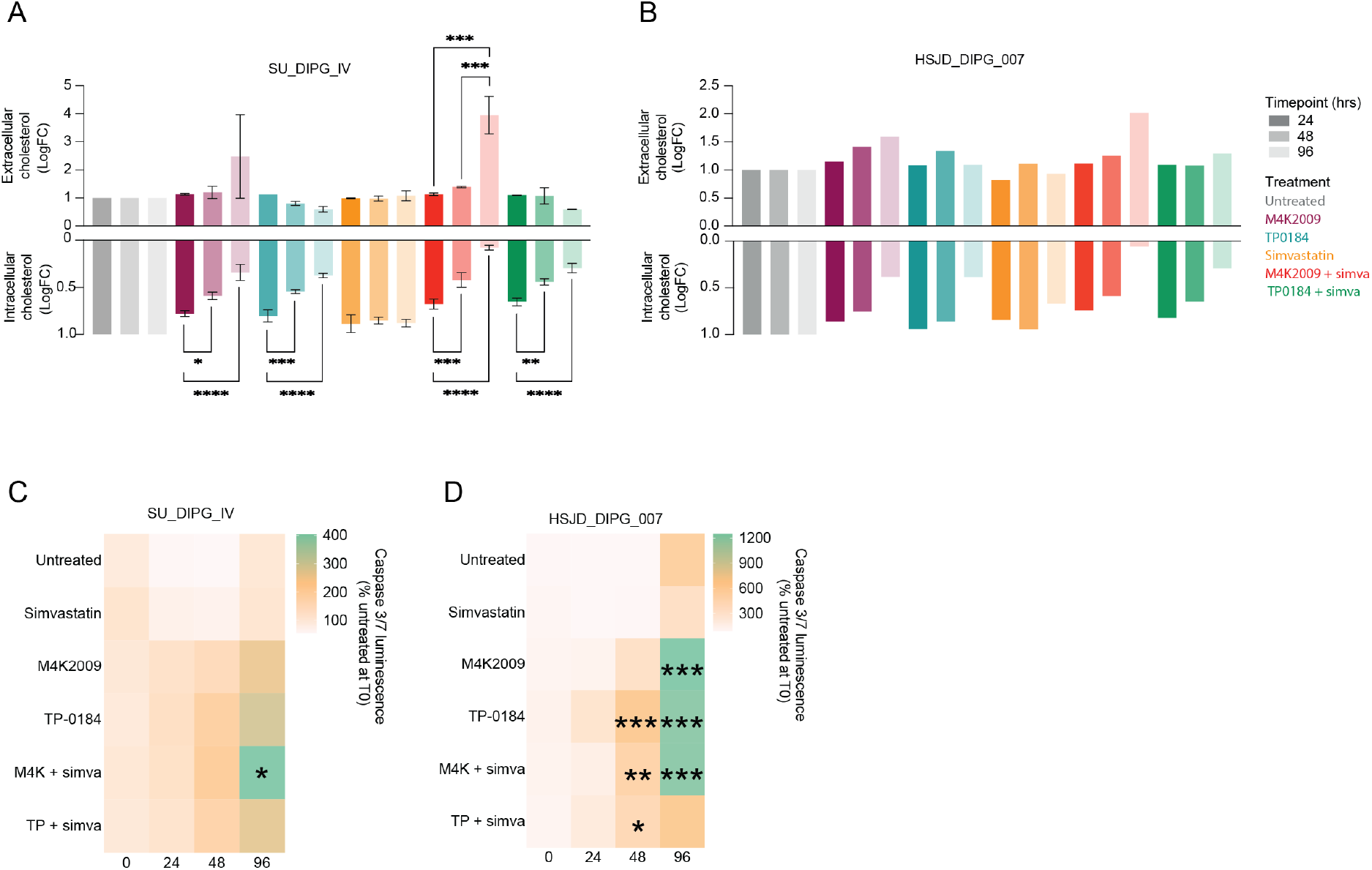
Time-course of intra- and extra-cellular cholesterol and caspase 3/7 activation following single-agent ALK2i or combination treatment. **A,** Log fold-change (LogFC, y-axis) of extracellular (top) and intracellular (bottom) Cholesterol-Glo luminescence following single-agent or combination treatment for 24, 48 or 96 h compared to untreated (DMSO) condition in SU_DIPG_IV cells. Data normalised to CellTiter-Glo and shown as mean ± range (n=2). **B,** Log fold-change (LogFC, y-axis) of extracellular (top) and intracellular (bottom) Cholesterol-Glo luminescence following single-agent or combination treatment for 24, 48 or 96 h compared to untreated (DMSO) condition in HSJD_DIPG_007. Data normalised to CellTiter-Glo (n=1). **C,** Heatmap of caspase 3/7 integrated luminescence intensity following single-agent or combination treatment for 24, 48 or 96 h compared to untreated (DMSO) condition in SU_DIPG_IV at time 0 (n=2). p values calculated by two-way ANOVA (* p<0.05, **p<0.01, ***p<0.001) compared to untreated condition at each time point. **D,** Heatmap of caspase 3/7 integrated luminescence intensity following single-agent or combination treatment for 24, 48 or 96 h compared to untreated (DMSO) condition in HSJD_DIPG_007 at time 0 (n=2). p values calculated by two-way ANOVA (* p<0.05, **p<0.01, ***p<0.001) compared to untreated condition at each time point.

**Extended data Fig.5:**
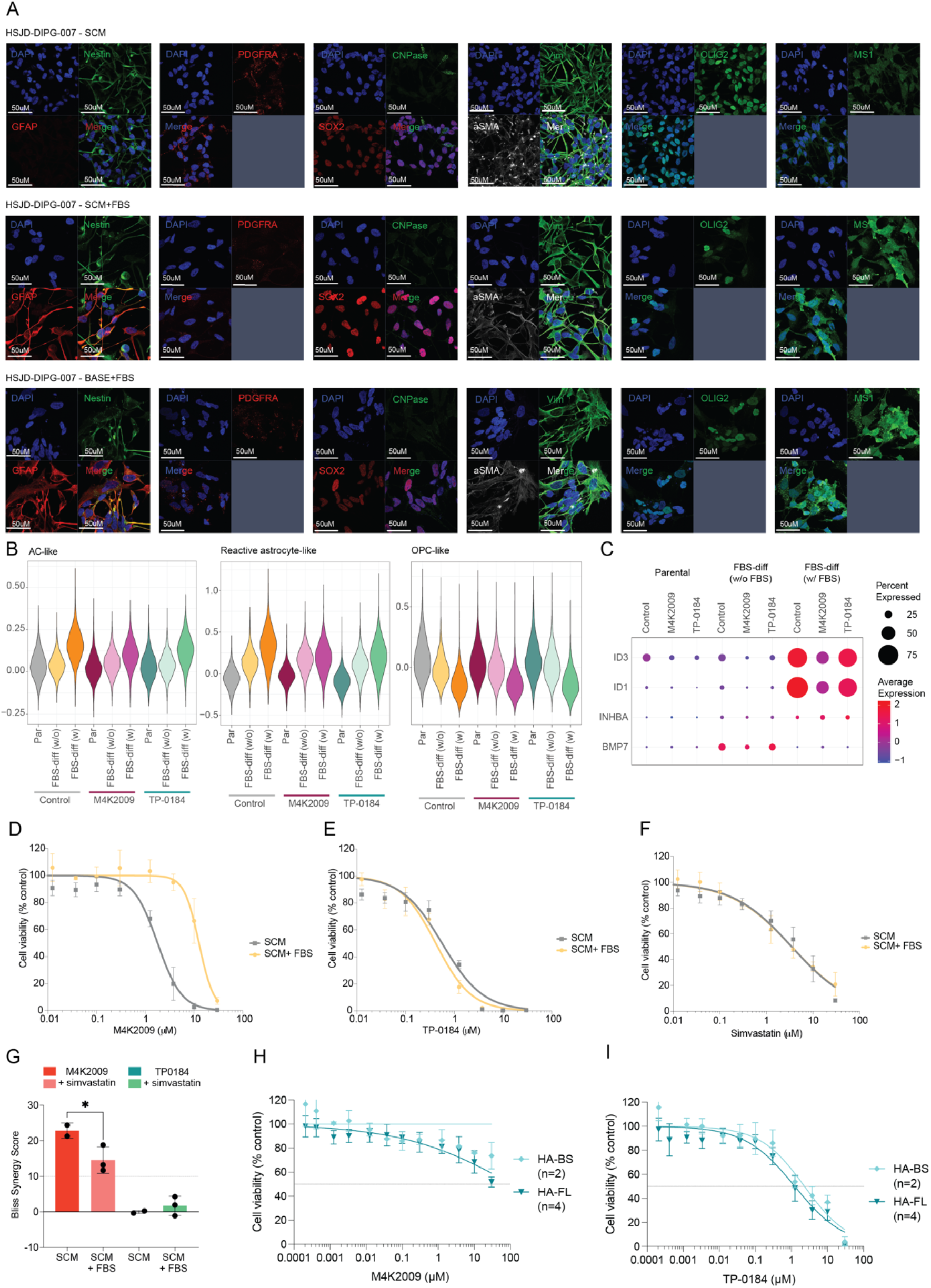
Characterisation of FBS-differentiated HSJD_DIPG_007 cultures. **A,** Representative images of nestin, MS1, SOX2, OLIG2, PDGFRA, GFAP, CNPase, aSMA and vimentin immunofluorescence staining in HSJD_DIPG_007 cells cultured in indicated media conditions. Images were acquired using the LSM700 at 40x magnification. Scale bar = 50μM. **B,** Violin plots showing the AC-like (left), reactive astrocyte-like (middle) and OPC-like (right) expression signature scores for each culture condition and drug treatment. The signature expression is plotted by colour according to the key provided. **C,** Bubble plot showing the single-cell RNAseq expression for *ID1*, *ID3*, *INHBA* (which encodes activin A) and *BMP7* in each culture condition with or without drug treatment. The average expression is plotted by colour according to the key provided. Circle size indicates the percentage of cells expressing the gene. **D,** Concentration-response curves for M4K2009 in HSJD_DIPG_007 cells cultured in SCM with or without 10 % FBS. Data normalised to DMSO vehicle control, shown as mean ± SD (n=3). Curves were fitted using Prism. **E,** Concentration-response curves for TP-0184 in HSJD_DIPG_007 cells cultured in SCM with or without 10 % FBS. Data normalised to DMSO vehicle control, shown as mean ± SD (n=3). Curves were fitted using Prism. **F,** Concentration-response curves for simvastatin in HSJD_DIPG_007 cells cultured in SCM with or without 10 % FBS. Data normalised to DMSO vehicle control, shown as mean ± SD (n=3). Curves were fitted using Prism. **G,** Bliss synergy scores for M4K2009 or TP-0184 plus simvastatin in HSJD_DIPG_007 cells cultured in SCM with or without 10 % FBS. Dashed line indicates Bliss synergy threshold (>10). Data shown as mean ± SD (n=3). p values calculated by one-way ANOVA (* p<0.05). **H,** Concentration-response curves for M4K2009 in normal human astrocytes isolated from the brainstem (HA-BS) or frontal-lobe (HS-FL). Data normalised to DMSO vehicle control, shown as mean ± SD (n≥2). Curves were fitted using [inhibitor] vs response, variable slope, four parameters in Prism. **I,**Concentration-response curves for TP-0184 in normal human astrocytes isolated from the brainstem (HA-BS) or frontal-lobe (HS-FL). Data normalised to DMSO vehicle control, shown as mean ± SD (n≥2). Curves were fitted using [inhibitor] vs response, variable slope, four parameters in Prism.

**Extended data Fig.6:**
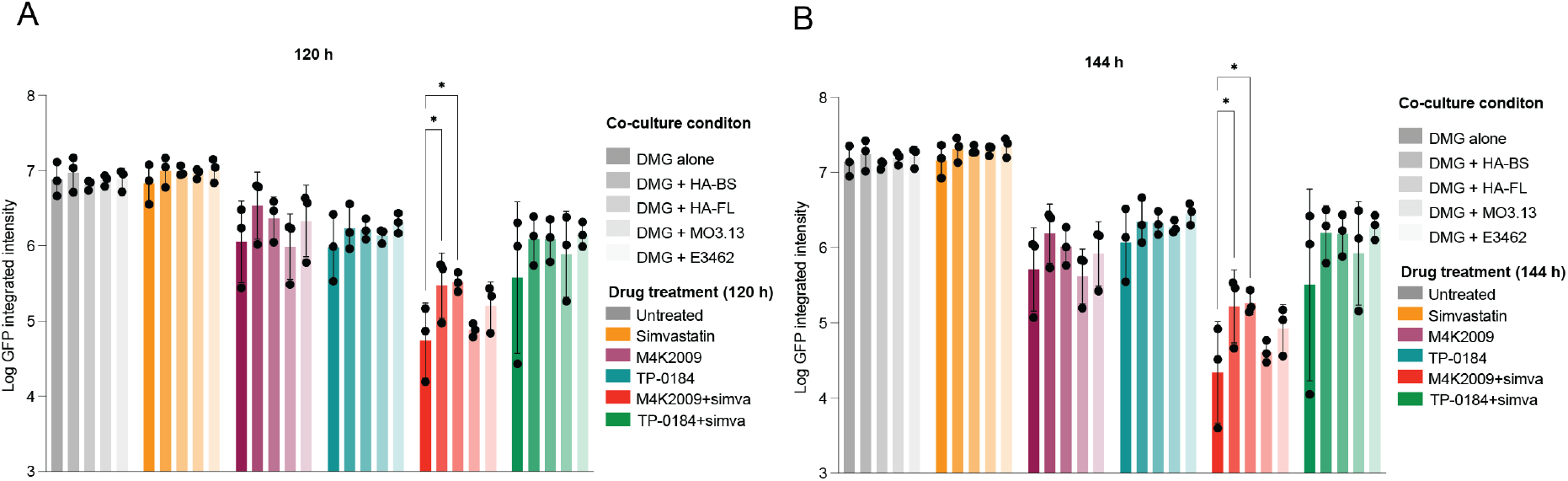
Assessing drug response of SU_DIPG_IV-GFP cells cultured alone or in co-culture. **A,** Log GFP integrated intensity for SU_DIPG_IV-GFP cultured alone or in co-culture following 120 h drug treatment. Data shown as mean ± SD (n=3). p values calculated by two-way ANOVA (* p<0.05). **B,** Log GFP integrated intensity for SU_DIPG_IV-GFP cultured alone or in co-culture following 144 h drug treatment. Data shown as mean ± SD (n=3). p values calculated by two-way ANOVA (* p<0.05).

